# Computational studies of drug repurposing targeting P-glycoprotein mediated multidrug-resistance phenotypes in agents of neglected tropical diseases

**DOI:** 10.1101/2020.06.12.147926

**Authors:** Nivedita Jaishankar, Sangeetha Muthamilselvan, Ashok Palaniappan

**Affiliations:** Department of Biotechnology, Sri Venkateswara College of Engineering, Post Bag No. 1, Pennalur, Sriperumbudur Tk 602117. India; Department of Bioinformatics, School of Chemical and BioTechnology, SASTRA Deemed University, Thanjavur 613401. India

**Keywords:** P-glycoprotein, neglected tropical diseases, multidrug resistance, homology modelling, receptor-ligand docking, differential ligand affinity, synergistic effects, leishmaniasis, trypanosomiasis, onchocerciasis, schistosomiasis

## Abstract

Mammalian ABCB_1_ P-glycoprotein is an ATP- dependent efflux pump with broad substrate specificity associated with cellular drug resistance. Homologous to this role in mammalian biology, the P-glycoprotein of agents of neglected tropical diseases (NTDs) mediates the emergence of multidrug-resistance phenotypes. The clinical and socioeconomic implications of NTDs are exacerbated by the lack of research interest among Big Pharma for treating such conditions. This work aims to characterise P-gp homologues in certain agents of key NTDs, namely

1. Protozoa: Leishmania major, Trypanosoma cruzi;
2. Helminths: Onchocerca volvulus, Schistosoma mansoni.

PSI-BLAST searches against the genome of each of these organisms confirmed the presence of P-gp homologues. Each homologue was aligned against five P-gp sequences of known structure, to identify the most suitable template based on sequence homology, phylogenetic nearest neighbor, and query coverage. Antibiotics used in the current line of therapy against each of these pathogens were identified using PubChem and their SMILES structures were converted to PDB using BABEL software. Potential antibiotics to test against the set of FDA-approved antibiotics were identified based on similarity to the chemical class of the known drugs and repurposing of the existing drugs. Docking studies of the respective modelled Pgp structures and the set of antibiotic ligands were carried out using AutoDock and the most tenable target-ligand conformations were assessed. The interacting residues within 4.5 Å of the ligand were identified, and the binding pockets were studied. The relative efficacy of the new drugs and the interacting pump residues were identified. Our studies could lay the foundation for the development of effective synergistic or new therapies against key neglected tropical diseases.

## 1. Introduction

### 1.1 Multidrug resistance (MDR)

Bacterial evolution has been constrained to respond to the selection pressure of antibiotics and combined with their reckless use, has led to the emergence of varied defenses against antimicrobial agents. The main mechanisms whereby the bacteria develop resistance to antimicrobial agents include enzymatic inactivation, modification of the drug target(s), and reduction of intracellular drug concentration by changes in membrane permeability or by the over expression of efflux pumps [1]. Multidrug resistance efflux pumps are recognized as an important component of resistance in both Gram-positive and Gram-negative bacteria [2]. Some bacterial efflux pumps may be selective for one substrate or transport antibiotics of different classes, conferring a multiple drug resistance (MDR) phenotype. With respect to efflux pumps, they provide a self-defense mechanism whereby antibiotics are extruded from the cell interior to the external environment. This results in sublethal drug concentrations at the active site that in turn may predispose the organism to the development of high-level target-based resistance [3]. Therefore, efflux pumps are viable antibacterial targets and identification and development of potent efflux pump inhibitors is a promising and valid strategy potential therapeutic agents that can rejuvenate the activity of antibiotics that are no longer effective against bacterial pathogens. The world is searching for new tools to combat multidrug resistance

### 1.2 P-GLYCOPROTEIN

P-glycoprotein is a mammalian multidrug resistance protein belonging to the ATP-binding cassette (ABC) superfamily (4). It is an ATP- dependent efflux pump encoded by the MDR1 gene and is primarily found in epithelial cells lining the colon, small intestine, pancreatic ductules, bile ductules, kidney proximal tubes, the adrenal gland, the blood-testis and the blood-brain barrier (5). This efflux activity of P-glycoprotein, coupled with its wide substrate specificity is responsible for the reduction in bioavailability of drugs as it extrudes all foreign substances such as drugs and xenobiotics out of the cells. ATP hydrolysis provides energy for the efflux of drugs from the inner leaflet of the cell membrane (6, 7). This protein is believed to have evolved as a defense mechanism against toxic compounds and prevent their entry into the cytosol (8).

P-glycoprotein confers resistance to a wide range of structurally and functionally diverse compounds, which has resulted in the emergence of multidrug resistance in medically relevant microorganisms. The pharmacodynamic role of P-glycoprotein in parasitic helminths has widespread clinical and socioeconomic implications, exacerbating the problem of neglected tropical diseases (NTDs) whose causative agents are helminths and protozoa.

Sheps et al., (9) reported that 15 P-glycoproteins are present in *Caenorhabditis elegans*, and Laing et al., (10) reported that 10 homologous P-glycoproteins were present in Haemonchus contortus. A bioinformatic and phylogenetic study conducted by Bourguinat et al., (11) on the Dirofilaria immitis genome identified three orthologous ABC-B transporter genes. These genes are suspected to be responsible for the P-glycoprotein mediated drug extrusion of melarsomine in D. immitus, and other parasites.

### 1.3 Neglected tropical diseases

Neglected Tropical Diseases (NTDs) encompass 17 bacterial, parasitic and viral diseases that prevail in tropical and subtropical conditions in 149 countries and affect more than 1 billion people worldwide (WHO – 2017).

#### 1.3.1 Leishmaniasis

Leishmaniasis, is a disease caused by parasites of the Leishmania type (WHO, 2014). It is spread by the bite of certain types of sandflies (12). The disease can present in three main ways: cutaneous, mucocutaneous, or visceral leishmaniasis (13). The cutaneous form presents with skin ulcers, while the mucocutaneous form presents with ulcers of the skin, mouth, and nose (12). Leishmaniasis is transmitted by the bite of infected female phlebotomine sandflies (14) which can transmit the protozoa *Leishmania*.

Gammaro et al., (12) first reported that the overexpression of P-glycoprotein in Leishmania species was responsible the drug resistance of the organisms against drugs such as methotrexate. The multidrug resistance has been associated with several ATP-binding cassette transporters including MRP1 (ABCC1) and P-glycoprotein (ABCB1). Wyllie et al., (15) demonstrated the presence of metal efflux pumps in the cell membrane of all Leishmania species. Soares et al., (16) reported that natural or synthetic modulators of human P-glycoprotein such as flavonoids, restore sensitivity to pentami-dine, sodium stibogluconate and miltefosine by modulating intracellular drug concentrations.

#### 1.3.2 Onchocerciasis

Onchocerciasis, also known as river blindness, is a disease caused by infection with the parasitic worm *Onchocerca volvulus* and is transmitted by the bite of an infected black fly of the *Simulium* type. . Symptoms include severe itching, bumps under the skin, and blindness. It is the second most common cause of blindness due to infection, after trachoma (WHO, 2014). Usually, many bites are required before infection occurs. A vaccine against the disease does not exist. Prevention is by avoiding being bitten by flies.

Ivermectin (IVM) is a semisynthesized macrocyclic lactone that belongs to the avermectin class of compounds. It is administered en mass and but is effective only against microfilariae (18). Bourguinat et al., (11) have found evidence of IVM resistance in Onchocerca volvulus. The clinical trial sampled patients before and after IVM treatment over a period of three years. The nodules collected from the patients contained IVM-resistant O. volvulus worms.

#### 1.3.3 Schistosomiasis

Schistosomiasis is a disease caused by infection with one of the species of *Schistosoma* helminthic flatworms known as flukes belonging to the class Trematoda of the phylum Platyhelminthes. There are three main species of Schistosoma associated with human disease: *Schistosoma mansoni* and *Schistosoma japonicum* cause intestinal schistosomiasis, and *Schistosoma haematobium* causes genitourinary schistosomiasis. Other Schistosoma species have been recognized less commonly as agents of intestinal schistosomiasis in humans (18). Pinto-Almeida *et al.,* (19) demonstrated that drug resistance by *Schistosoma mansoni* to praziquantel (commonly employed drug) is mediated by efflux pump proteins, including P-glycoprotein and multidrug resistance-associated proteins.

#### 1.3.4 Trypanosomiasis

The trypanosomiasis consists of a group of diseases caused by parasitic protozoa of the genus Trypanosoma. There are two main parasites such as *Trypanosoma brucei* which causes the sleeping sickness or human African trypanosomiasis and *Trypanosoma cruzi* which causes the Chagas’ disease or American trypanosomiasis (20). These diseases are transmitted by several arthropod vectors such as Glossina and Triatomine. Chaga’s disease causes 21,000 deaths per year mainly in Latin America (21). Benznidazole and Nifortimox only available drugs however have limited efficacy in the advanced stages of the disease (22). Liu et al., (23) and Rappa et al., (24) concluded that Trypanosoma cruzi develops resistance to the drugs after prolonged treatment. It was shown that this happens due to the overexpression of the MDR1 gene, at high levels of the drug, which accumulates in the cells over time. Campos et al., (25) demonstrated that the drug resistance is continued throughout the life cycle of the worm.

## 2. Methods

The methodology is essentially similar to that in our earlier study on P-glycoproteins in priority infectious agents (26).

### 2.1 Determining the full helminthic complement of efflux pump proteins homologous to mammalian p-glycoprotein

The protein sequence of the human P-glycoprotein (P08183) was obtained from the SWISS-PROT database. The position-specific iterated BLAST (PSI-BLAST) was performed against a search set of non-redundant protein sequences in the organism of interest, using hPGP as the query. Through a PSI-BLAST search, a large set of related proteins are compiled. It is used to identify distant evolutionary relationships between protein sequences (27). In The algorithm parameters were set with an E-value of 0.001, and the scoring matrix BLOSUM62 was used. This step was performed on all four organisms of interest (*Leishmania major, Onchocerca volvulus, Schistosoma mansoni* and *Trypanosoma cruzi*). Hundreds of hits were obtained for P-glycoprotein, and these results were prioritized according to pre-determined parameters such as medical relevance, annotation status and the presence of conserved regions. Sequences having a high percentage of sequence identity and query coverage were prioritized. Specific UniProt searches of these protein sequences were performed using the Accession number. The results were analyzed, and the P-glycoprotein sequence of each organism was finalized.

### 2.2 Multiple Sequence Alignment

The templates chosen for multiple sequence alignment (MSA) were 4M1M (*Mus musculus*), 4F4C (*Canorhabditis elegans*), 3WME (*Cyanidioschyzon merolae*), 2HYD (*Staphylococcus aureus*), 3B5Z (*Salmonella enterica*). These five metazoan, algal and bacterial templates were used due to their high sequence identity with the hPGP sequence (Palaniappan et al., 2016). The target sequences and the five templates were aligned using ClustalX 2.1 (28). MSA was performed in order to infer the homology and evolutionary relationship between the sequences of the biological dataset. The clustering algorithm used was Neighbour Joining (NJ). The phylogenetic distance between the target sequence and the templates was calculated.

### 2.3 Homology Modelling

The chosen p-glycoprotein sequences were used as target sequences for modelling using software such as SWISS-MODEL. SWISS-MODEL is an open-source, structural bioinformatics tool used for the automated comparative modeling of three-dimensional protein structures (29, 30). Several P-glycoprotein structures were modelled for each organism, using multiple templates. The templates having high sequence similarity with the target sequences were given preference. The models were built, and the PDB files of the structures were obtained.

### 2.4 Structure validation

The validity of the structures was checked using Procheck, an open-source tool used to assess the reliability of the protein structure. It is a part of the SWISS-MODEL server. The structures were refined using energy-minimisation protocols and the least energetic structure corresponding to each protein was chosen for docking studies. The criteria used to assess the quality of the structure include model geometry and the Ramachandran plot. The Ramachandran plot describes the rotation of the polypeptide backbone around the N-C_α_ (◻) and C-C_α_ (ψ) bonds. It provides an overview of the distribution of the torsion angles over the core, allowed, generous and disallowed regions. The three main parameters used to select the structures were:

1. Overall Ramachandran value
2. Phylogenetic tree distance
3. Taxonomy

### 2.5 Creation of the Ligand Dataset

The ligand dataset was created by surveying the literature to determine the drugs which the pathogenic helminths are both sensitive and resistant to. Drug resistance which was conferred via efflux pump activity was given importance. This set of ligands was created for each efflux pump, comprising of known and potential antibiotics. The canonical SMILES (*simplified molecular-input line-entry system*) of each drug was retrieved from the PubChem database. The PDB model of each antibiotic was then generated using MarvinView by converting the canonical SMILES (31).

### 2.6 Protein and Ligand Preparation

The efflux pump proteins and ligands were individually docked using the AutoDock Version 4.2.6 suite of programs (32). The software consists of two main programs: AutoGrid, which pre-calculates a set of grid points on the receptor, and AutoDock, which docks the ligand to the receptor through the grids. The PDB files of the P-glycoprotein structures and the ligands were modified through the addition of Gasteiger charges, followed by the addition and merging of hydrogen atoms to each structure. These modified structures were then saved as PDBQT files using the AutoDock tools. A uniform grid box was then defined and centered in the internal binding cavity of each P-glycoprotein structure, and the affinity maps were generated using AutoGrid. This procedure was repeated for each protein-drug complex.

### 2.7 Molecular Docking of the Helminthic Efflux Pumps with Known and Potential Antibiotics

Each drug was individually docked with each target protein using AutoDock 4.2.6. The local search algorithm used was the Lamarckian genetic algorithm, set to its default parameters. The docking parameters were set to 250000 cycles per run and 10 runs per protein-drug complex, to obtain the 10 best poses for each complex. The best pose was defined as the conformation having the least binding energy. The ten poses obtained for each receptor-ligand pair were clustered at 2.0 *Å* r.m.s. to validate the convergence to the best pose. The AutoDock was run, and the PDBQT file of the best pose of each docked complex was generated.

The results were analysed to verify whether the pathogenic strain could develop resistance to known antibiotics using efflux pump activity and if the novel antibiotics could be effective against the development of such resistance.

### 2.8 Calculation of Differential Ligand Binding Affinity

The differential binding affinities of the repurposed ligands were determined using the conventionally used drugs as a baseline. A lower value is indicative of a more stable complex. The differential affinity of the potential drug for a given efflux pump protein relative to the known drug is estimated as the difference in the binding energies of the known and potential drugs.

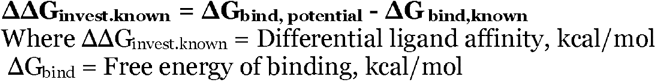

### 2.9 Identification of Interacting Residues in Each Docked Complex

The best pose of each docked complex was viewed using RasMol (33) and all interacting residues within a radius of 4.5 *Å* of the ligand were selected. The PDBQT file of each restricted complex was saved as a PDB file. The interacting residues of each docked complex were then analyzed.

## 3. Results and Discussion

Extensive literature searches on Neglected Tropical Diseases (NTDs) showed that leishmaniasis, onchocerciasis, schistosomiasis and trypanosomiasis have started exhibiting multidrug resistance, mediated by P-glycoprotein efflux pumps (11, 12, 25, 34). New drugs targeting NTD’s are undergoing clinical trials (35–37) and efforts are being taken to uncover the mechanisms of drug resistance employed by the causative helminths.

The sequence identity of each helminthic P-glycoprotein with the human P-glycoprotein (hPGP) which was retrieved from the UniProt database (UniProt ID: P08183) was determined by running a PSI-BLAST.

### 3.1 PSI-BLAST ANALYSIS

The PSI-BLAST was performed on each target organism using hPGP as the query. The results were refined according to pre-determined parameters such as medical relevance, annotation status and the presence of conserved regions. The chosen efflux pump protein sequences were shown in table 1.

**Table 1:** PSI-BLAST results of the target organisms using hPGP as the query.

| Organism | Name of protein | Sequence length | % of identity | Query coverage | Max score |
| --- | --- | --- | --- | --- | --- |
| *Leishmania major* | p-glycoprotein | 1341 | 36% | 98% | 767 |
| *Onchocerca volvulus* | p-glycoprotein | 1278 | 37% | 97% | 776 |
| *Schistosoma mansoni* | SMDR2 | 1254 | 40% | 98% | 889 |
| *Trypanosoma cruzi* | p-glycoprotein | 1034 | 29% | 30% | 79.7 |

The top hits of each PSI-BLAST were analyzed, and the hit having the highest Max Score was chosen only in the case of *Leishmania major* and *Onchocerca volvulus*. These protein sequences were fully annotated and had high sequence identities over a large portion of the protein sequence. The top hits of the PSI-BLAST of *Schistosoma mansoni* and *Trypanosoma cruzi* with hPGP yielded results having high Max Scores, but low query coverage. These protein sequences were also found to be unannotated. For these reasons, the proteins which had a lower Max Score in comparison to other results, but satisfied other parameters, were chosen.

### 3.2 Template Selection and Multiple Sequence Alignment

The following metazoan, algal and bacterial crystal structures were selected as potential templates for homology modeling (38). This justifies their use as templates for MSA in all subsequent steps.

Each target protein sequence was aligned with the chosen template using ClustalX 2.1. The MSA between *Leishmania major* and the 4M1M and 4F4C templates showed the highest sequence identity, as shown in Figure 1. Thus, these two templates were given the highest priority in all succeeding steps.

**Figure 1:**
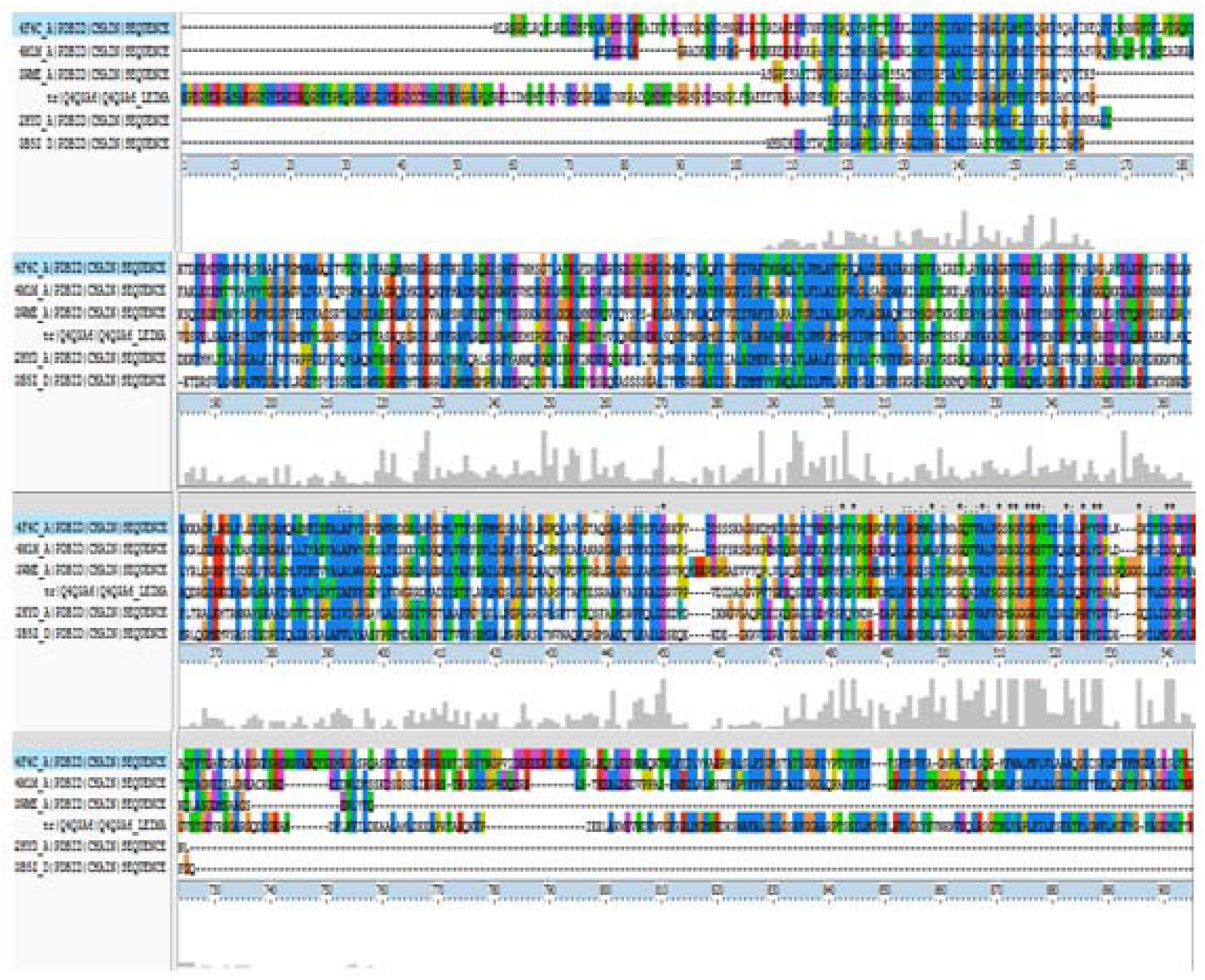
Multiple sequence alignment of P-glycoprotein [*Leishmania major*] (target sequence) with the templates of interest. Identical residues are marked with *.

Additionally, the MSA was performed and the phylogenetic distance between the organisms was calculated (shown in Table. 3). The clustering algorithm used was NJ.

**Table 2:** Templates chosen for multiple sequence alignment.

| TEMPLATE | ORGANISM |
| --- | --- |
| 4M1M | *Mus musculus* |
| 4F4C | *Canorhabditis elegans* |
| 3WME | *Cyanidioschyzon merolae* |
| 2HYD | *Staphylococcus aureus* |
| 3B5Z | *Salmonella enteric* |

**Table 3:** Phylogenetic distance between the target sequence of each organism and each template.

| Template | PHYLOGENETIC DISTANCE |  |  |  |
| --- | --- | --- | --- | --- |
|  | *Leishmania major* | *Onchocerca volvulus* | *Schistosoma mansoni* | *Trypanosoma cruzi* |
| **4M1M** | 0.685 | 0.648 | 0.646 | **0.847\*** |
| **4F4C** | **0.642** | **0.638** | **0.605** | 0.861 |
| **3WME** | 0.653 | 0.679 | 0.649 | 0.867 |
| **2HYD** | 0.72 | 0.709 | 0.694 | 0.826 |
| **3B5Z** | 0.731 | 0.707 | 0.698 | 0.841 |

*The distance between T. cruzi and 4M1M is prioritized as the 4M1M and 4F4C templates were found to have a higher sequence identity with the helminths.

### 3.3 HOMOLOGY MODELLING

The chosen P-glycoprotein sequences of the organisms were used as target sequences for homology modeling using the SWISS-MODELLER. Each protein was modeled using several templates, and the pre-determined templates were used if they were found to have a fairly high GMQE score. Each modeled structure was saved as a PDB file. The results are summarized in Table 4.

**Table 4:** Homology modeling results.

| Organism | Template | Sequence identity | Query coverage | GMQE |
| --- | --- | --- | --- | --- |
| ***Leishmania major*** | 4F4C | 34.43 | 0.91 | 0.64 |
|  | 4M1M | 36.09 | 0.90 | 0.65 |
|  | 3WME | 37.30 | 0.43 | 0.29 |
|  | 4Q9I | 36.20 | 0.90 | 0.65 |
|  | 4KSC | 38.25 | 0.90 | 0.64 |
|  | 4KSB | 38.25 | 0.90 | 0.64 |
|  | 5KPJ | 38.25 | 0.90 | 0.66 |
| ***Onchocerca volvulus*** | 4F4C | 38.83 | 0.97 | 0.69 |
|  | 4M1M | 37.10 | 0.96 | 0.69 |
|  | 3WME | 31.75 | 0.44 | 0.33 |
|  | 3G5U | 38.77 | 0.95 | 0.66 |
|  | 4KSB | 38.77 | 0.95 | 0.67 |
|  | 4Q9I | 36.97 | 0.95 | 0.69 |
|  | 4LSG | 38.77 | 0.95 | 0.67 |
| ***Schistosoma mansoni*** | 4F4C | 38.60 | 0.97 | 0.69 |
|  | 4M1M | 36.09 | 0.90 | 0.65 |
|  | 3G5U | 39.41 | 0.97 | 0.68 |
|  | 3G60 | 42.11 | 0.95 | 0.68 |
|  | 5KPJ | 39.52 | 0.97 | 0.70 |
|  | 4KSC | 42.11 | 0.95 | 0.69 |
|  | 4KSB | 42.11 | 0.95 | 0.69 |
|  | 4LSG | 39.41 | 0.97 | 0.69 |
| ***Trypanosoma cruzi*** | 4F4C | 15.11 | 0.81 | 0.45 |
|  | 4M1M | 14.36 | 0.78 | 0.29 |
|  | 3WME | 18.37 | 0.51 | 0.33 |
|  | 4KSC | 14.23 | 0.79 | 0.44 |
|  | 3G5U | 14.46 | 0.78 | 0.43 |
|  | 5TSI | 23.43 | 0.90 | 0.52 |
|  | 4LSG | 14.46 | 0.78 | 0.43 |

GMQE (Global Model Quality Estimation) is a score which provides an estimation of the quality of the alignment. It is expressed as a value between 0 and 1, where the reliability of the model is directly proportional to the score. The GMQE of the homology models are found to be (mostly) between 0.60 and 0.70 for all organisms, with the exception of *Trypanosoma cruzi* which gave scores in the range 0.29 to 0.52.

The templates 4M1M, 4F4C and 3WME were found to be comparatively more reliable. Hence, only the protein structures modeled using these templates were used for further validation studies.

### 3.4 STRUCTURE VALIDATION

The quality of each structure was assessed using Procheck. Criteria such as model geometry and the Ramachandran plot were used to validate the structures. The PDB file of each structure was used to run the Procheck, and the Ramachandran plot values were obtained. The Ramachandran values are summarized in Table 5.

**Table 5:** Justification of the template chosen for each organism using the Ramachandran plot values and the phylogenetic distance between the target protein and the template.

| *Leishmania major* |  |  |  |  |  |  |  |  |
| --- | --- | --- | --- | --- | --- | --- | --- | --- |
| Template | Core region | Additionally allowed region | Generously allowed region | Disallowed region | No. of residues | Query coverage | Sequence identity | Phylogenetic distance |
| 4F4C | 90.8 | 8.1 | 1.2 | 0 | 1250 | 0.43 | 34.43 | 0.685 |
| 4M1M | 92 | 6.4 | 1.4 | 0.2 | 1188 | 0.9 | 36.09 | 0.642 |
| 3WME | 94.6 | 4.6 | 0.4 | 0.4 | 573 | 0.43 | 36.46 | 0.653 |
| *Onchocerca volvulus* |  |  |  |  |  |  |  |  |
| Templat | Core region | Additionally allowed region | Generously allowed region | Disallowed region | No. of residues | Query coverage | Sequence identity | Phylogenetic distance |
| 4F4C | 90.8 | 8.1 | 1.2 | 0 | 1250 | 0.97 | 38.83 | 0.648 |
| 4M1M | 91.1 | 7.6 | 1.2 | 0.1 | 571 | 0.95 | 37.1 | 0.638 |
| 3WME | 93.1 | 5.8 | 0.6 | 0.6 | 1180 | 0.45 | 33.51 | 0.679 |
| *Schistosoma mansoni* |  |  |  |  |  |  |  |  |
| Templat | Core region | Additionally allowed region | Generously allowed region | Disallowed region | No. of residues | Query coverage | Sequence identity | Phylogenetic distance |
| 4F4C | 90.8 | 8.1 | 1.2 | 0 | 1250 | 0.97 | 38.6 | 0.646 |
| 4M1M | 91.1 | 6.9 | 1.8 | 0.2 | 572 | 0.46 | 36.6 | 0.605 |
| 3WME | 93.7 | 5.8 | 0.2 | 0.4 | 567 | 0.45 | 38.31 | 0.649 |
| *Trypanosoma cruzi* |  |  |  |  |  |  |  |  |
| Templat | Core region | Additionally allowed region | Generously allowed region | Disallowed region | No. of residues | Query coverage | Sequence identity | Phylogenetic distance |
| 4F4C | 90.8 | 8.1 | 1.2 | 0 | 1250 | 0.81 | 15.11 | 0.847 |
| 4M1M | 86.0 | 11.9 | 1.8 | 0.2 | 1034 | 0.51 | 17.80 | 0.867 |
| 3WME | 89.8 | 8 | 1.6 | 0.6 | 573 | 0.51 | 18.37 | 0.861 |

The structures were finalized by analyzing overall Ramachandran value, Phylogenetic tree distance, Taxonomy parameters. The 4F4C template was found to suitable for all the organisms excluding *Leishmania major*, for which the 4M1M template was, selected (Table 5).

#### 3.4.1 Validation of the P-glycoprotein structure modeled using the 4M1M template for Leishmania major

The Ramachandran plots having a core region of at least 90% are prioritized for further studies. The core, allowed, generous and disallowed regions are coloured and distinguished (Figure 2). The red, brown, and yellow regions represent the favored, allowed, and generously allowed regions.

**Figure 2:**
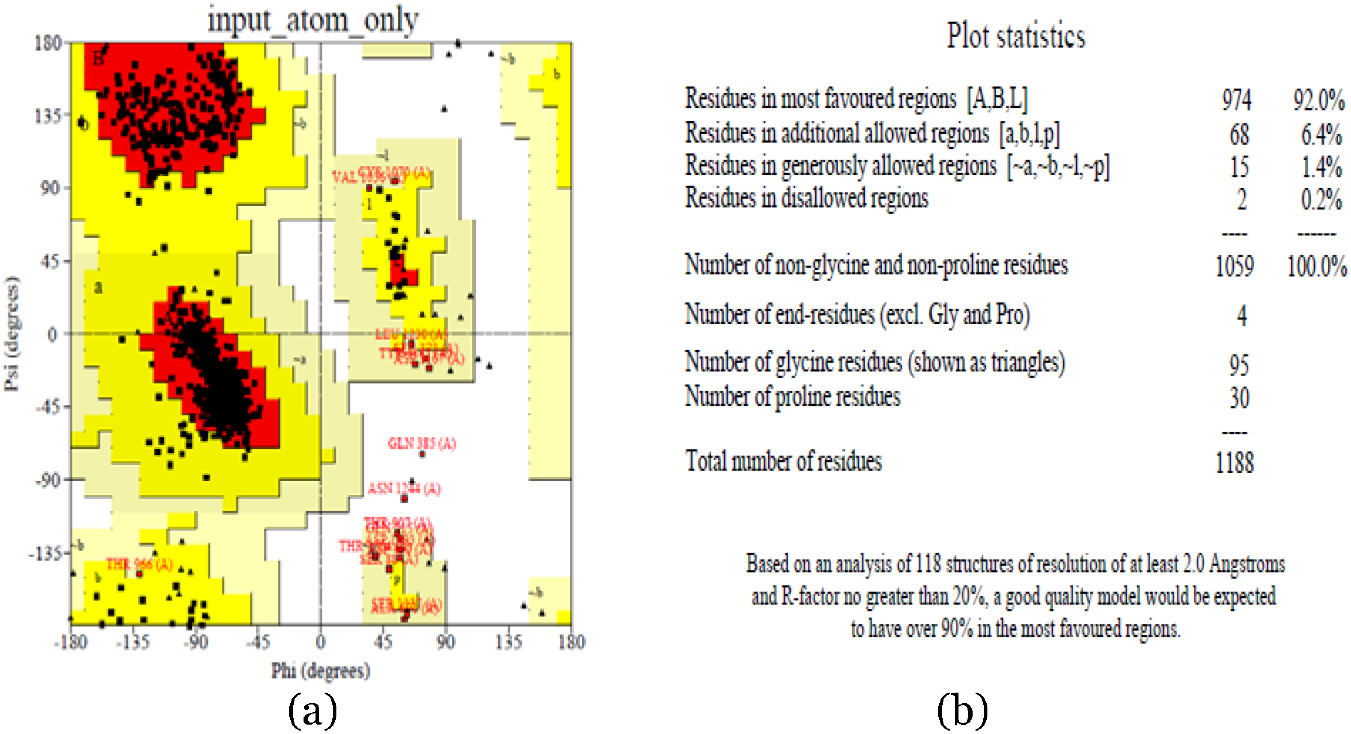
**(a)** The Ramachandran plot generated for P-glycoprotein [*Leishmania major*], modeled using the 4M1M template **(b)** Plot statistics of the P-glycoprotein [*Leishmania major*], modeled using the 4M1M template.

A more comprehensive analysis of the structure is provided by other programs which generate other data such as Phi- Psi graphs and Chi1-Chi2 plots for each residue type.Each Phi- Psi plot provides an analysis of the torsion angle of each residue type. The red, brown, and yellow regions represent the favored, allowed, and generously allowed regions (shown in Figure 3).

**Figure 3:**
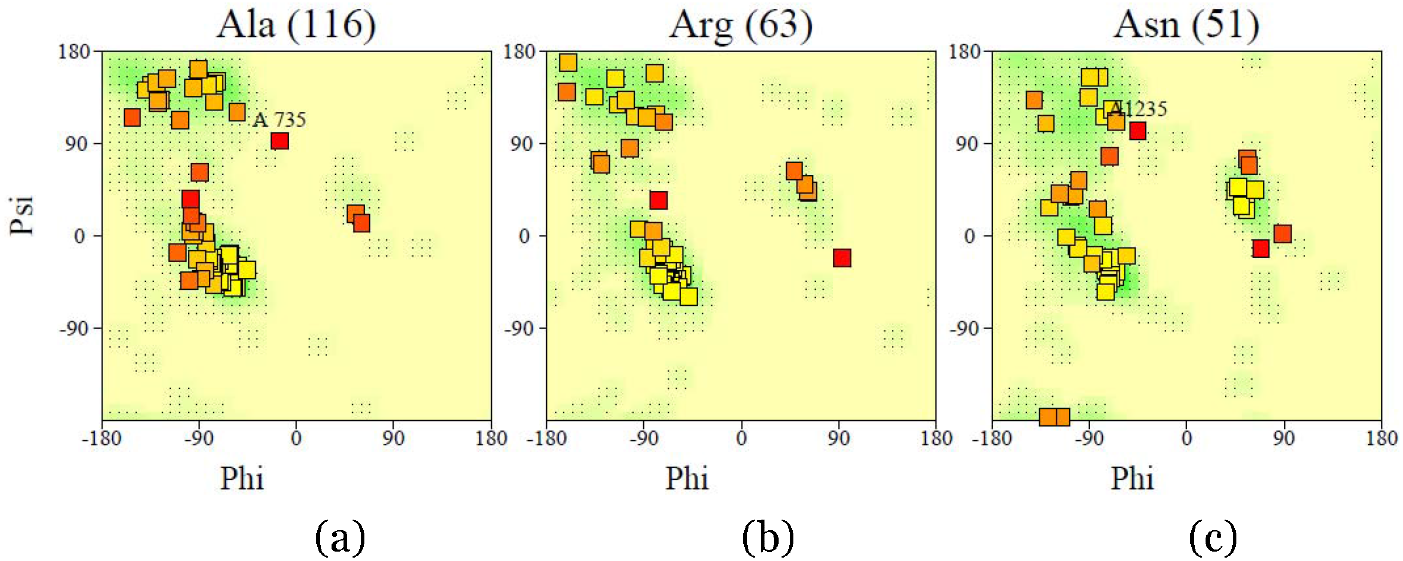
Phi- Psi plot of residues of the P-glycoprotein structure of *Leishmania major*, modeled using the 4M1M template (a) Ala, (b) Arg and (c) Asn

The Chi1-Chi2 plot describes the side-chain torsion angles combinations for each amino acid (28). The darker regions indicate a more favourable angle combination (shown Figure 4).

**Figure 4:**
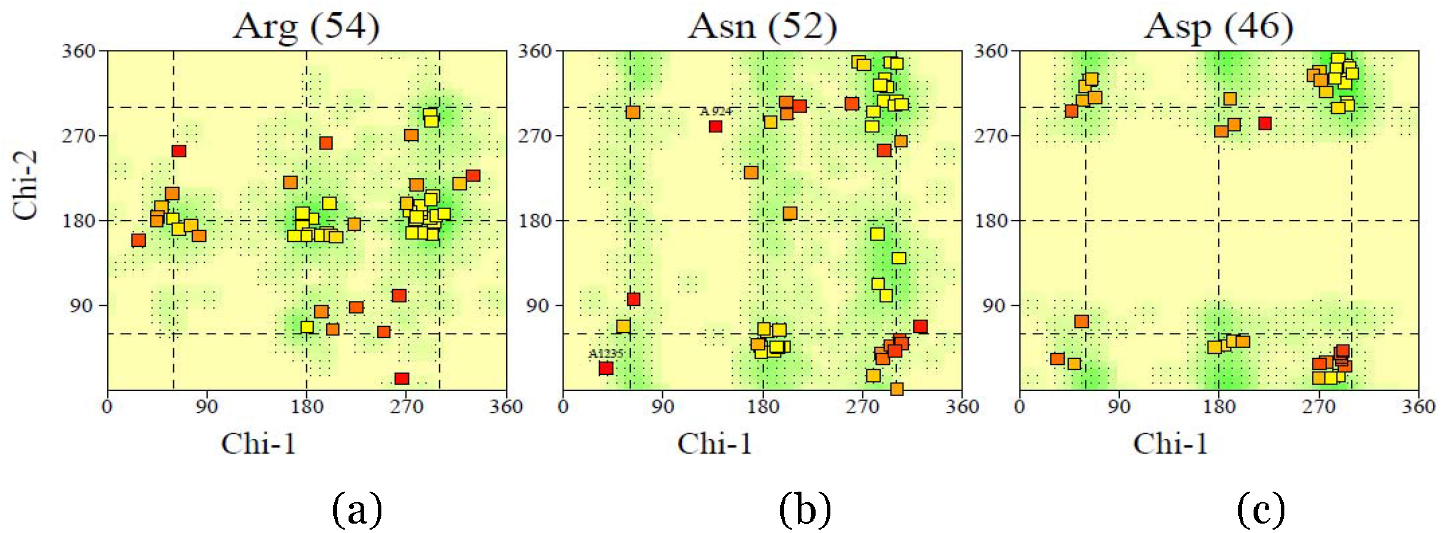
Chi1- Chi2 plot of residues of the P-glycoprotein structure of *Leishmania major*, modeled using the 4M1M template (a)Arg, (b)Asn and (c)Asp

#### 3.4.2 Validation of the P-glycoprotein structure modeled using the 4F4C template for *Onchocerca volvulus*

The Ramachandran plot obtained for this P-glycoprotein structure, modeled using the 4F4C template shows a core region value of 90.8% (Figure 5). Figure 6 provides an analysis of the torsion angle of each residue type. The darker regions indicate a more favourable angle combination (shown Figure 7)

**Figure 5:**
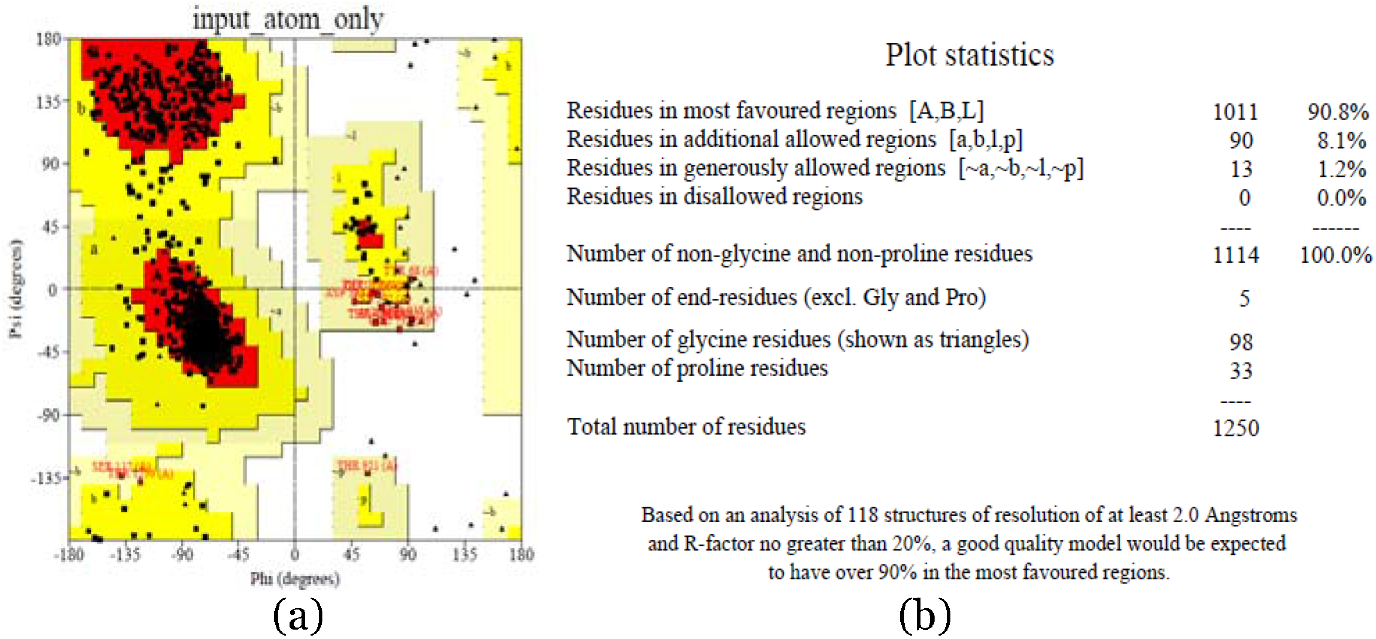
**(a)** The Ramachandran plot generated for P-glycoprotein [*Onchocerca volvulus*], modeled using the 4F4C template **(b)** Plot statistics of the P-glycoprotein [*Onchocerca volvulus*], modeled using the 4F4C template.

**Figure 6:**
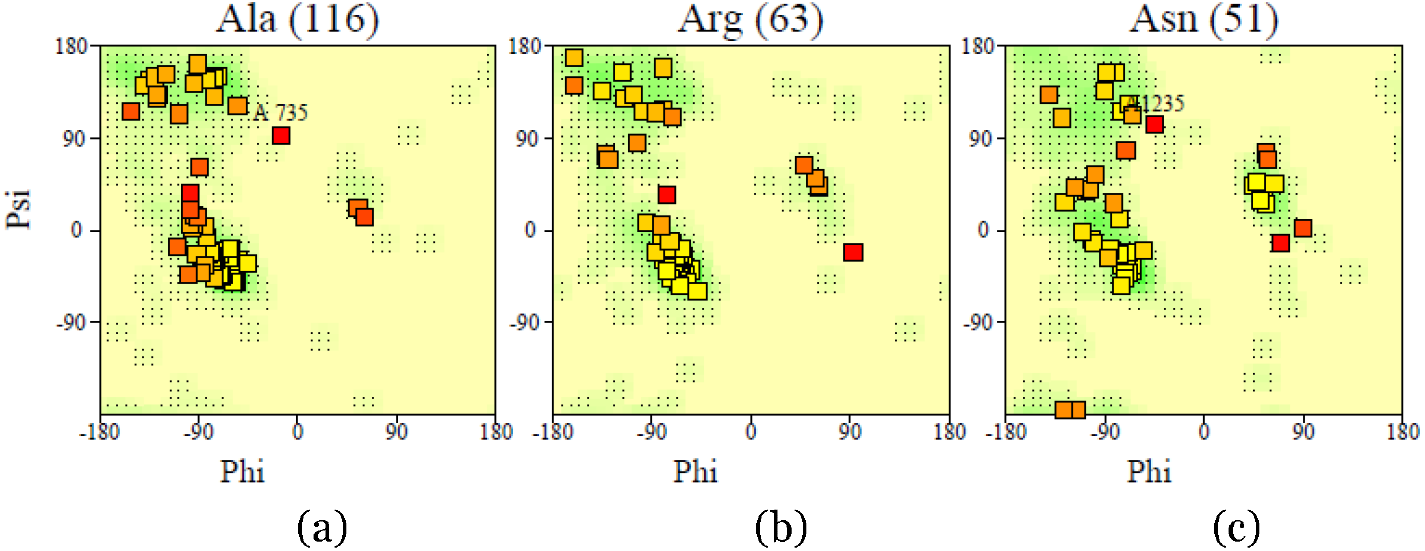
Phi- Psi plot of residues of the P-glycoprotein structure of *Onchocerca volvulus*, modeled using the 4F4C template (a) Ala, (b)Arg and (c)Asn

**Figure 7:**
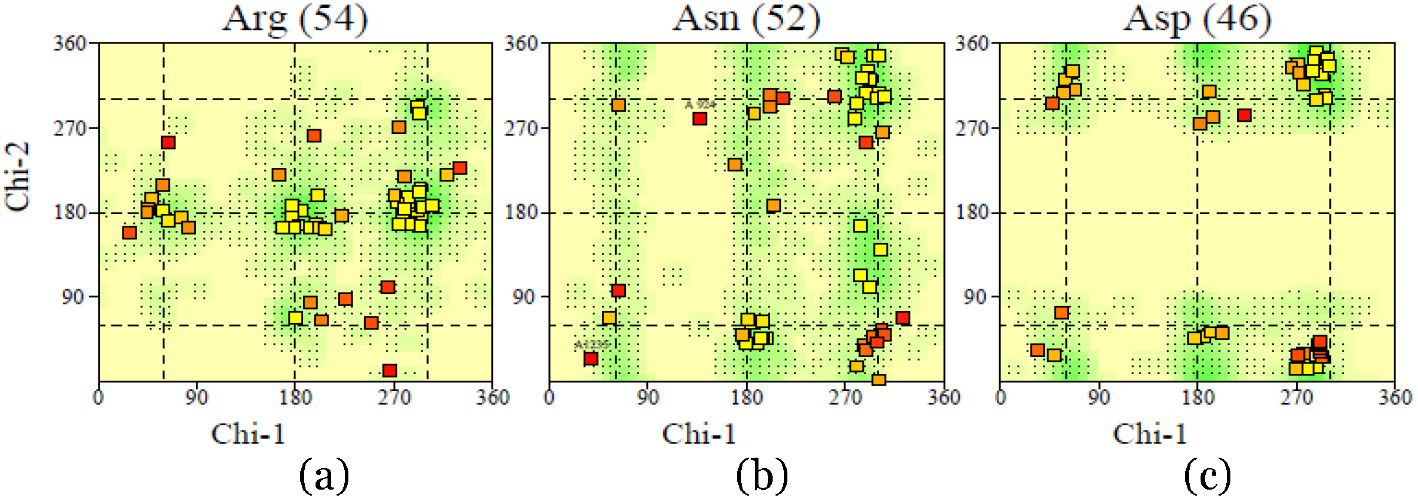
Chi1- Chi2 plot of residues of the P-glycoprotein structure of *Onchocerca volvulus*, modeled using the 4F4C template. (a) Arg,(b) Asn and (c)Asp

#### 3.4.3 Validation of the P-glycoprotein structure modeled using the 4F4C template for *Schistosoma mansoni*

The Ramachandran plot obtained for this P-glycoprotein structure, modeled using the 4F4C template shows a core region value of 90.8% (Figure 8). Figure 9 provides an analysis of the torsion angle of each residue type. The darker regions indicate a more favourable angle combination (shown Figure 10)

**Figure 8:**
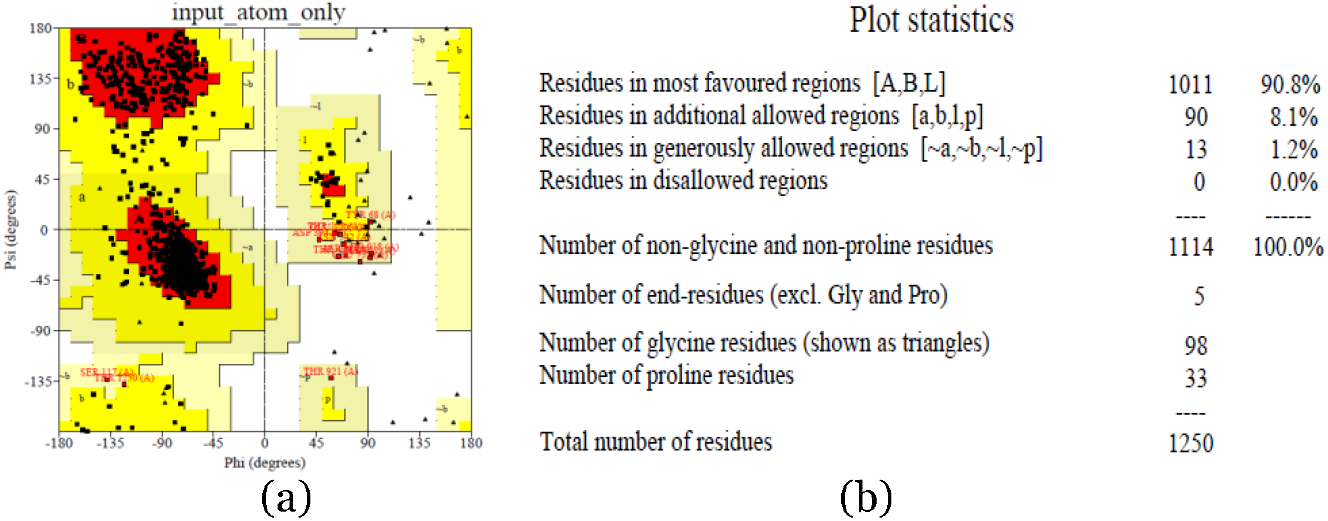
**(a)** The Ramachandran plot generated for P-glycoprotein [*Schistosoma mansoni*], modeled using the 4F4C template **(b)** Plot statistics of the P-glycoprotein [*Schistosoma mansoni*], modeled using the 4F4C template.

**Figure 9:**
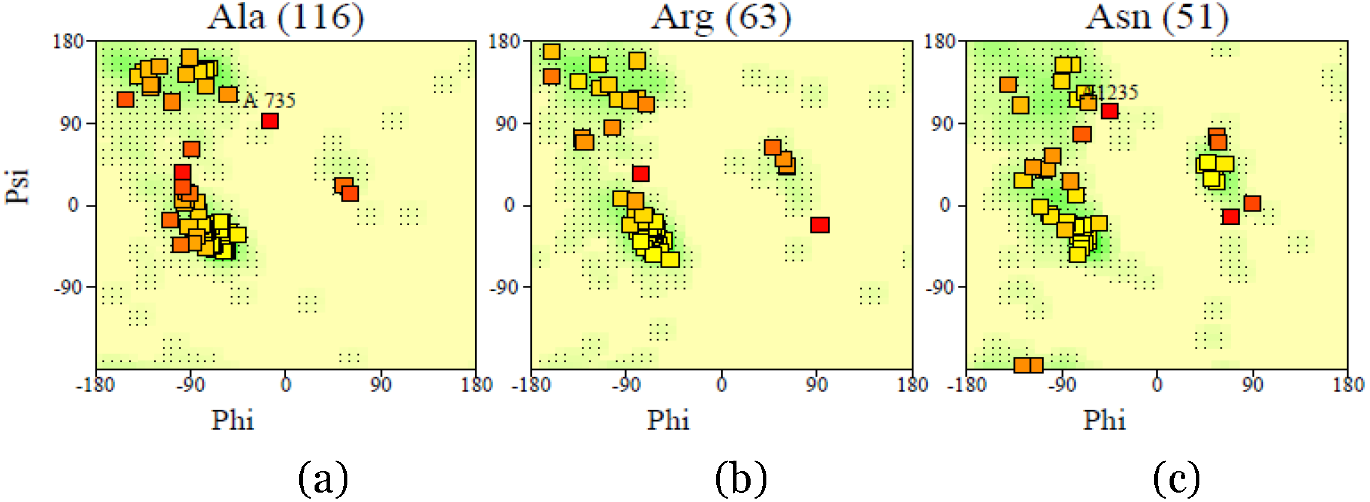
Phi- Psi plot of residues of the P-glycoprotein structure of *Schistosoma mansoni*, modeled using the 4F4C template. (a) Ala, (b) Arg and (c) Asn

**Figure 10:**
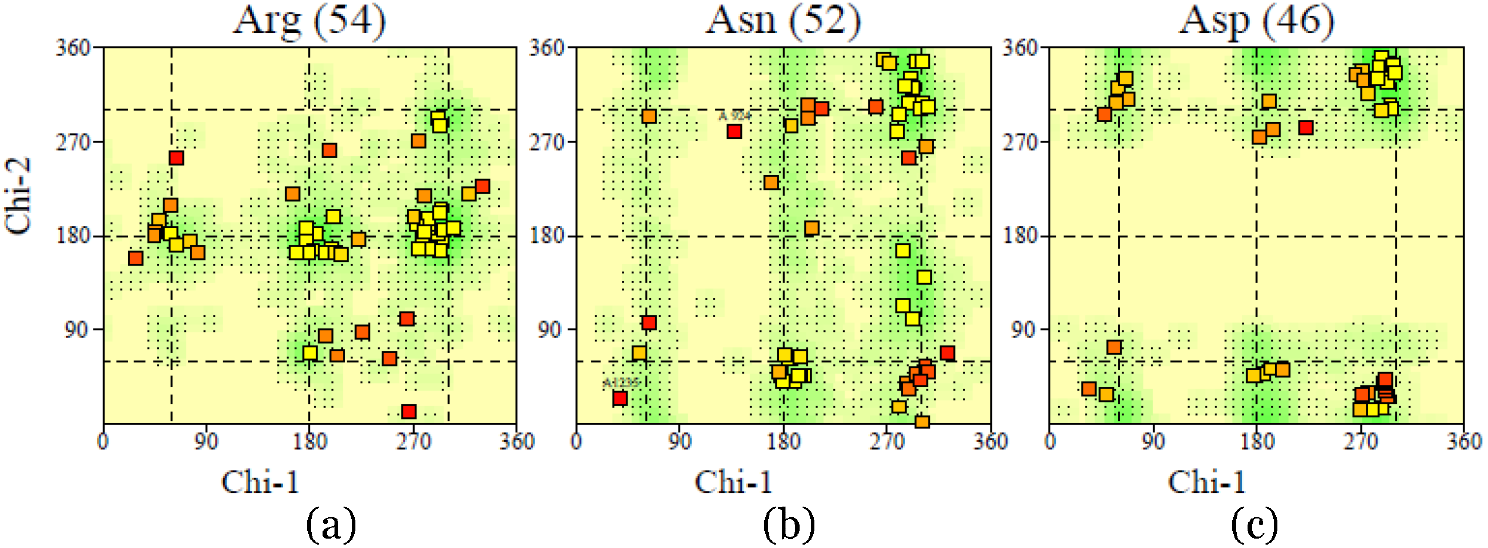
Chi1- Chi2 plot of residues of the P-glycoprotein structure of *Schistosoma mansoni*, modeled using the 4F4C template (a) Arg, (b) Asn and (c) Asp

#### 3.4.4 Validation of the P-glycoprotein structure modeled using the 4F4C template for *Trypanosoma cruzi*

The Ramachandran plot obtained for this P-glycoprotein structure, modeled using the 4F4C template shows a core region value of 90.8% (11). Figure 12 provides an analysis of the torsion angle of each residue type. The darker regions indicate a more favourable angle combination (shown Figure 13)

**Figure 11:**
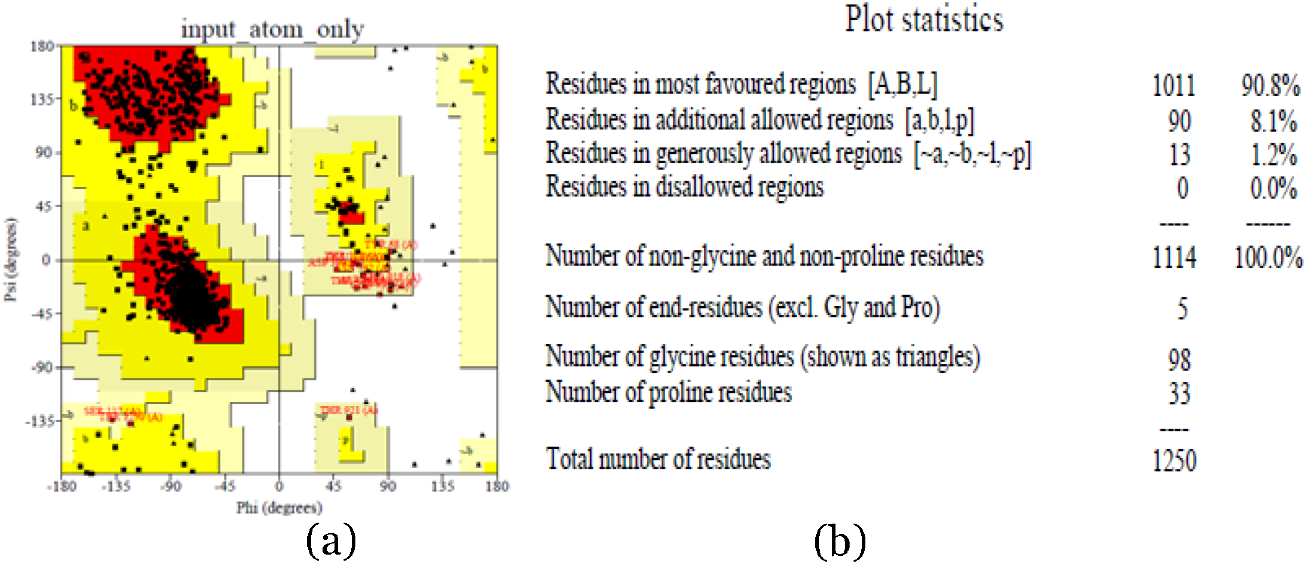
**(a)** The Ramachandran plot generated for P-glycoprotein [*Trypanosoma cruzi*], modeled using the 4F4C template **(b)** Plot statistics of the P-glycoprotein [*Trypanosoma cruzi*], modeled using the 4F4C template.

**Figure 12:**
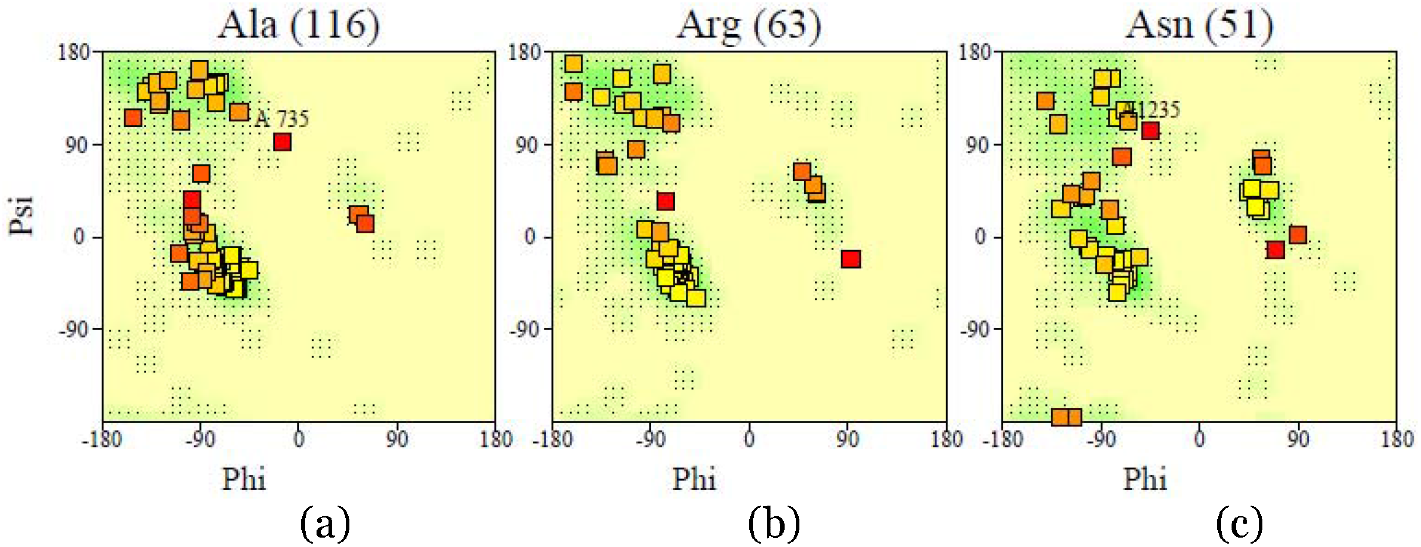
Phi- Psi plot of residues of the P-glycoprotein structure of *Trypanosoma cruzi*, modeled using the 4F4C template (a) Ala, (b) Arg and (c) Asn

**Figure 13:**
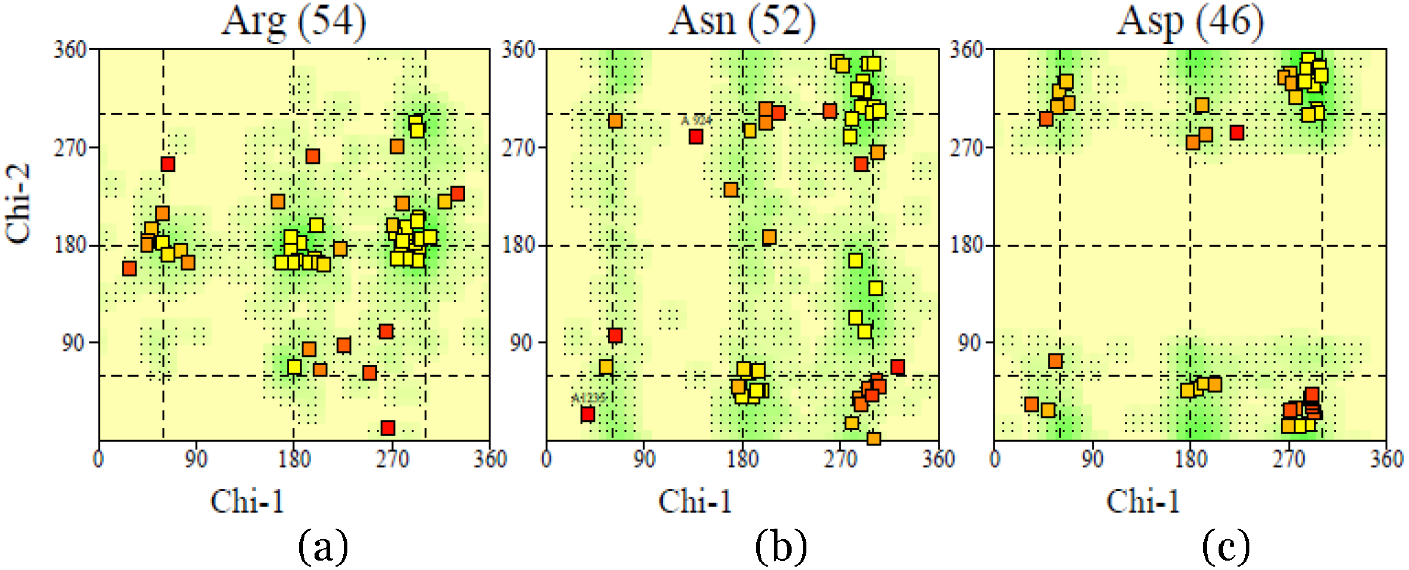
Chi1- Chi2 plot of residues of the P-glycoprotein structure of *Trypanosoma cruzi*, modeled using the 4F4C template (a) Arg, (b) Asn and(c) Asp

### 5.5 CREATION OF THE LIGAND DATASET

Upon extensive survey of the literature, a comprehensive dataset of the known and potential drugs was compiled. The list of potential drugs comprises of both unapproved, investigational drugs which are undergoing phase trials, and FDA approved antibiotics. In this study, these known drugs have been repurposed for other helminthic diseases.

### 3.6 MOLECULAR DOCKING OF THE HELMINTHIC EFFLUX PUMPS WITH KNOWN AND POTENTIAL ANTIBIOTICS

The molecular docking was carried out using the AutoDock suite of tools. The search algorithm used was the Lamarckian Genetic Algorithm, and the docking parameters were set to 10 runs per protein-drug complex. Each docked complex yielded 10 poses, and the best pose was defined as the conformation possessing the least free binding energy.

#### 3.6.1 Molecular docking results of benznidazole with P-glycoprotein [*Leishmania major*]

The drug benznidazole is docked with P-glycoprotein [*Leishmania major*], and their interaction is studied (Table 7). The best pose has a free binding energy of −5.00 kcal/mol. The clustering was performed at 2.0 *Å* r.m.s. to validate the convergence to the best pose. The clustering figure (Figure 14) shows closer peaks near −2.5 kcal/mol, whereas the least binding energy of the complex, i.e. most clustering is at −5.66 kcal/mol. This shows that convergence to the best pose can be achieved through consecutive dockings with more iterations. Figure 14(b) depicts the binding site on the receptor and figure 15(c) shows the interacting residues in the benznidazole- P-glycoprotein [*Leishmania major*]) docked complex viewed through RasMol 2.1

**Table 6:**
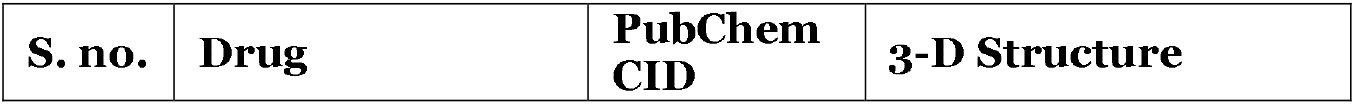
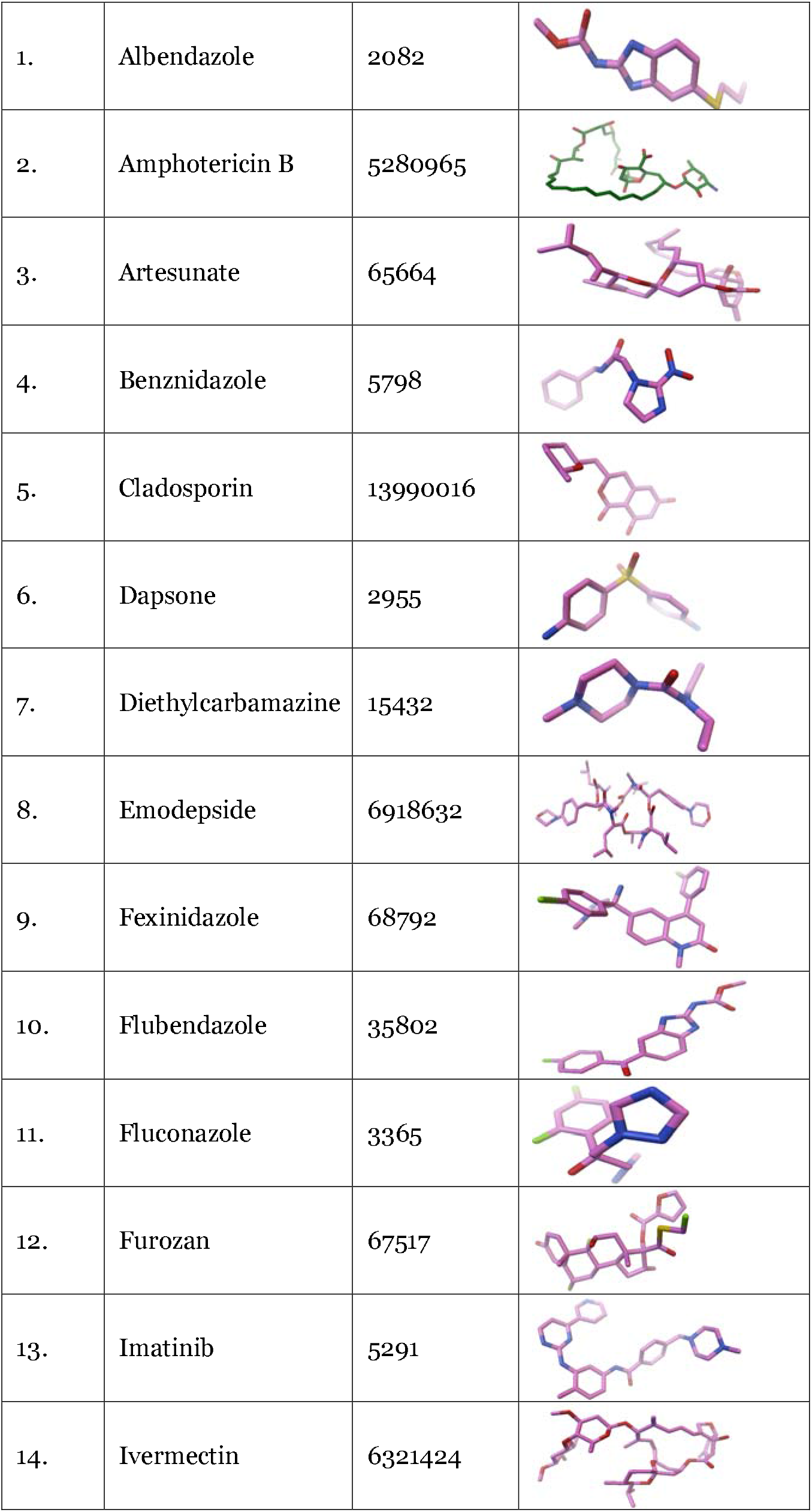
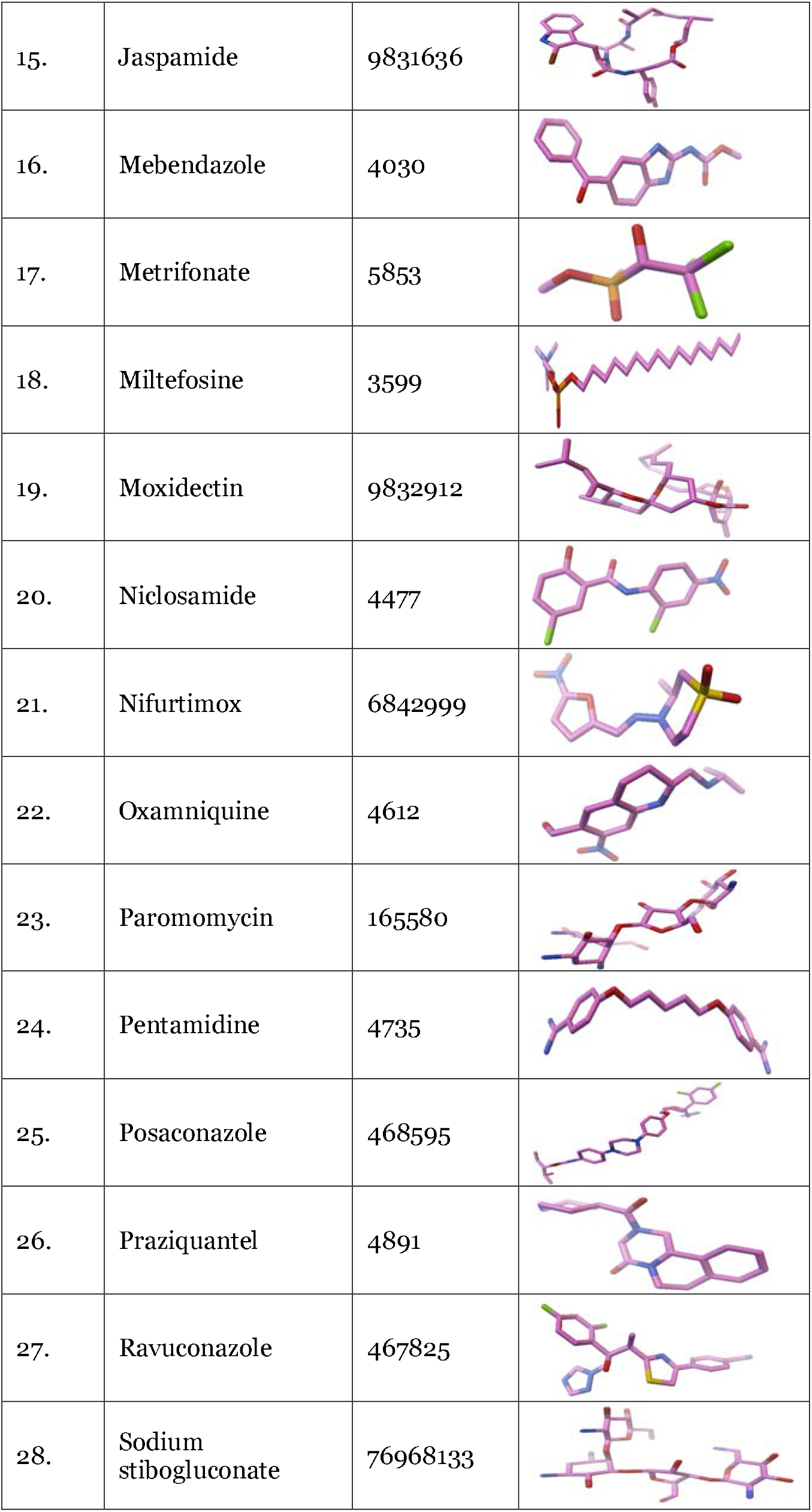
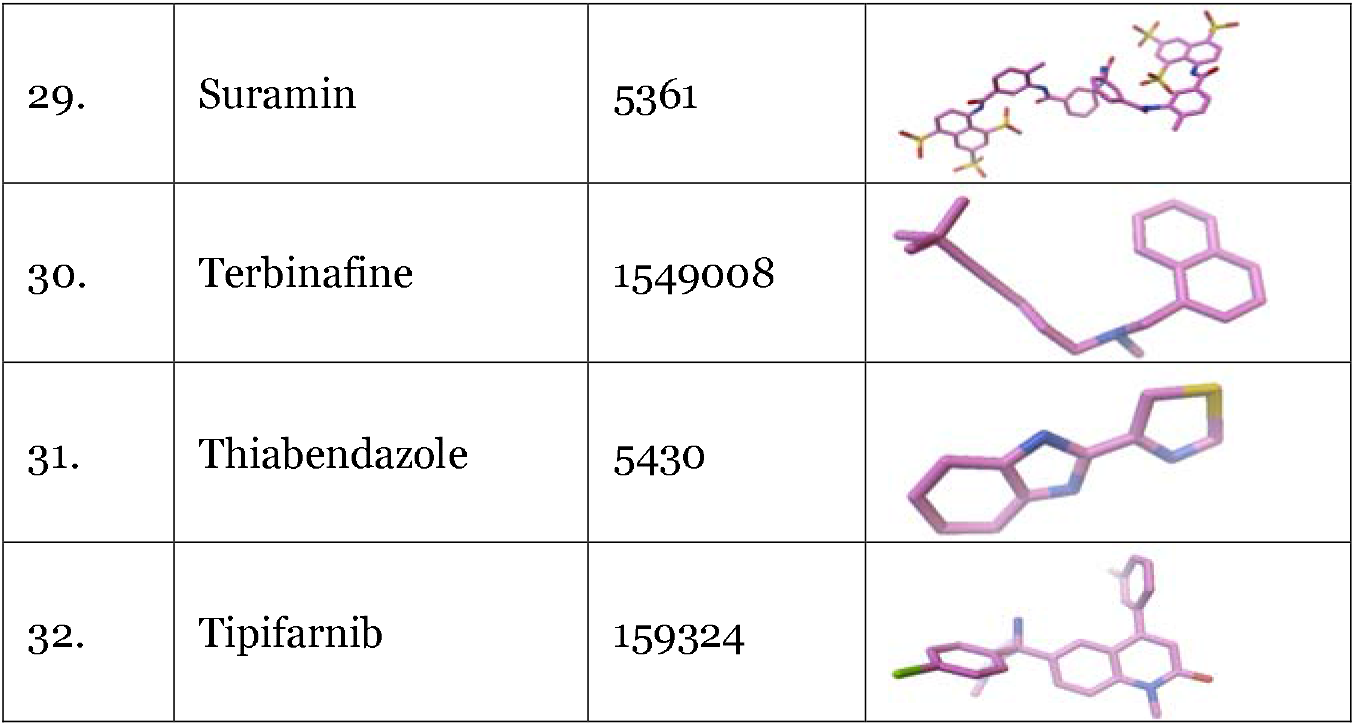
PubChem Compound ID and 3-D structure of the ligands used for docking studies. 3-D structures of the drugs are visualized using Python Molecular Viewer (PMV-1.5.6).

| S. no. | Drug | PubChem CID | 3-D Structure |
| --- | --- | --- | --- |
| 1. | Albendazole | 2082 |  |
| 2. | Amphotericin B | 5280965 |  |
| 3. | Artesunate | 65664 |  |
| 4. | Benznidazole | 5798 |  |
| 5. | Cladosporin | 13990016 |  |
| 6. | Dapsone | 2955 |  |
| 7. | Diethylcarbamazine | 15432 |  |
| 8. | Emodepside | 6918632 |  |
| 9. | Fexinidazole | 68792 |  |
| 10. | Flubendazole | 35802 |  |
| 11. | Fluconazole | 3365 |  |
| 12. | Furozan | 67517 |  |
| 13. | Imatinib | 5291 |  |
| 14. | Ivermectin | 6321424 |  |
| 15. | Jaspamide | 9831636 |  |
| 16. | Mebendazole | 4030 |  |
| 17. | Metrifonate | 5853 |  |
| 18. | Miltefosine | 3599 |  |
| 19. | Moxidectin | 9832912 |  |
| 20. | Niclosamide | 4477 |  |
| 21. | Nifurtimox | 6842999 |  |
| 22. | Oxamniquine | 4612 |  |
| 23. | Paromomycin | 165580 |  |
| 24. | Pentamidine | 4735 |  |
| 25. | Posaconazole | 468595 |  |
| 26. | Praziquantel | 4891 |  |
| 27. | Ravuconazole | 467825 |  |
| 28. | Sodium stibogluconate | 76968133 |  |
| 29. | Suramin | 5361 |  |
| 30. | Terbinafine | 1549008 |  |
| 31. | Thiabendazole | 5430 |  |
| 32. | Tipifarnib | 159324 |  |

**Table 7:** Interaction of the drug benznidazole with P-glycoprotein [*Leishmania major*]

| Rank of Complex | Free Binding Energy (kcal/mol) |
| --- | --- |
| 1 | -5.00 |
| 2 | -4.84 |
| 3 | -4.2 |
| 4 | -4.41 |
| 5 | -3.77 |
| 6 | -3.48 |
| 7 | -2.96 |
| 8 | -2.64 |
| 9 | -2.54 |
| 10 | -2.48 |

**Figure 14:**
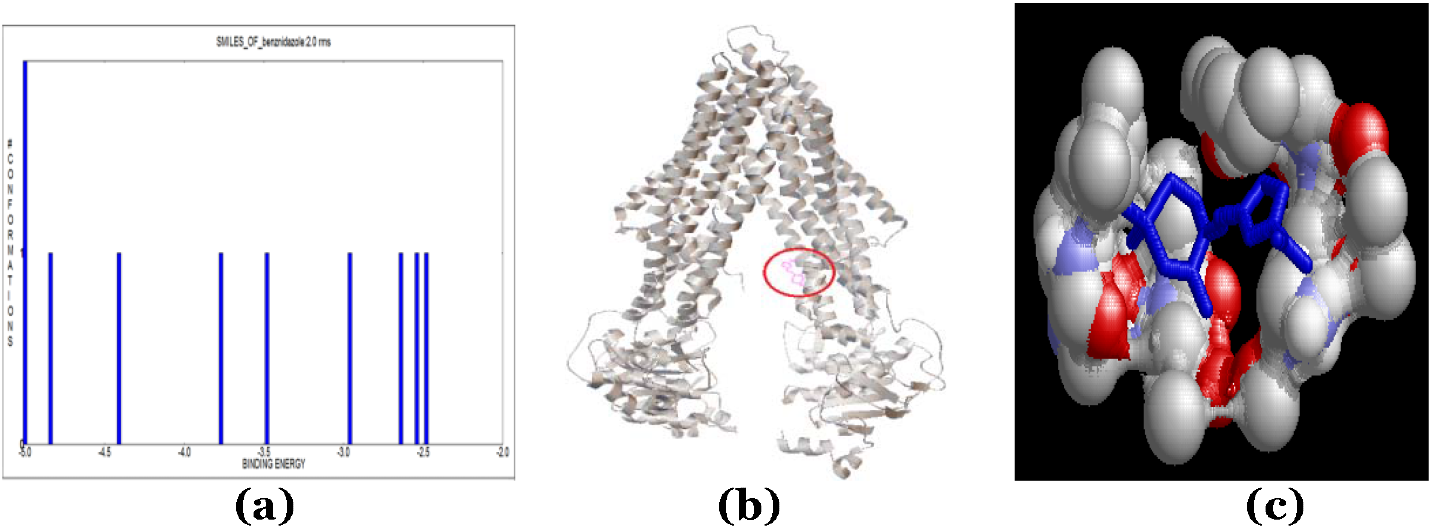
(a)Clustering analysis of the benznidazole- P-glycoprotein docked complex. (b) Location of the binding site on the receptor (P-glycoprotein [Leishmania major]). (c) The interacting residues in the benznidazole- P-glycoprotein [*Leishmania major*]) docked complex is viewed using RasMol 2.1.

**Figure 15:**
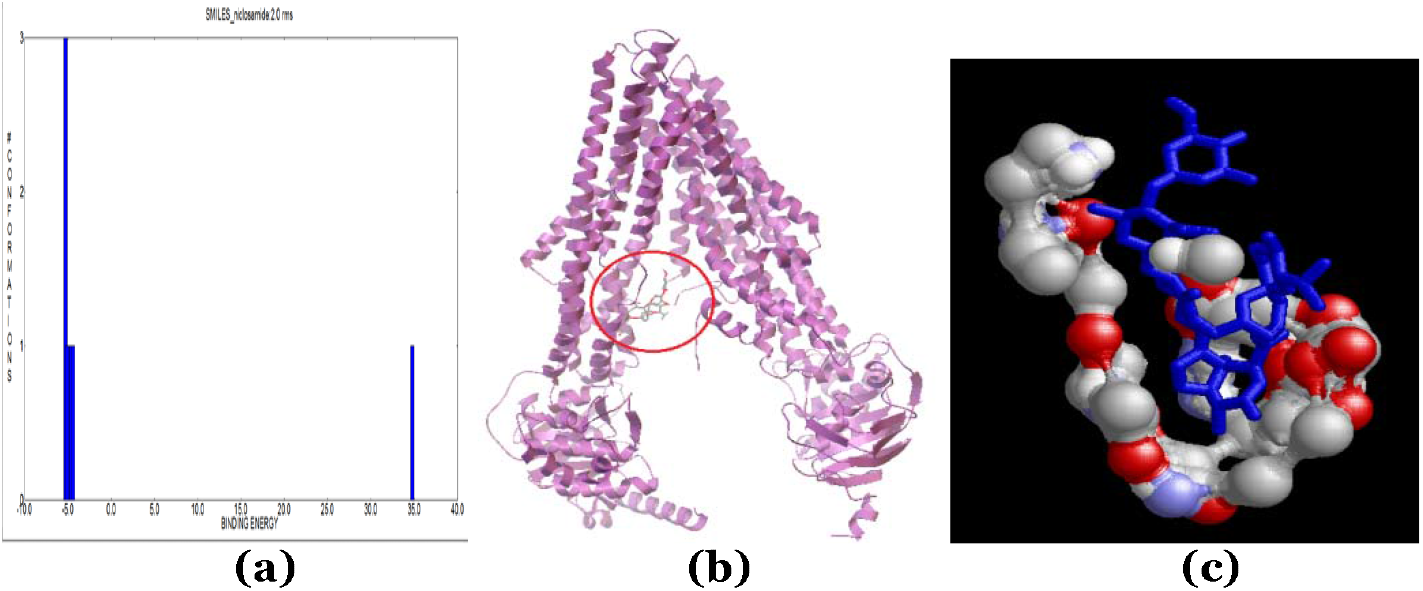
(a) Clustering analysis of the niclosamide-P-glycoprotein docked complex. (b) Location of the binding site on the receptor (P-glycoprotein *Onchocerca volvulus*]). (c) The interacting residues in the niclosamide-P-glycoprotein [*Onchocerca volvulus*]) docked complex is viewed using RasMol 2.1.

#### 3.6.2 Molecular docking results of niclosamide with P-glycoprotein [*Onchocerca volvulus*]

The best pose has a free binding energy of −5.29 kcal/mol (Table 8). The clustering figure shows the most number of conformations at −1.30 kcal/mol (Figure 15). Figure 15(b) depicts the binding site on the receptor and figure 15(c) shows the interacting residues in the niclosamide-P-glycoprotein [*Onchocerca volvulus*]) docked complex viewed through RasMol 2.1

**Table 8:** Interaction of the drug niclosamide with P-glycoprotein [*Onchocerca volvulus*]

| Rank of Complex | Free binding energy (kcal/mol) |
| --- | --- |
| 1 | -5.29 |
| 2 | -5.01 |
| 3 | -4.78 |
| 4 | -5.14 |
| 5 | -5.08 |
| 6 | -5.02 |
| 7 | -4.59 |
| 8 | -4.53 |
| 9 | -4.42 |
| 10 | 34.78 |

#### 3.6.3 Molecular docking results of praziquantel with P-glycoprotein [*Schistosoma mansoni*]

The best pose has a free binding energy of −5.83 kcal/mol (Table 9). The clustering figure (Figure 16) shows the most number of conformations at −5.0 kcal/mol. Figure 16(b) depicts the binding site on the receptor and figure 22 shows the interacting residues in the Praziquantel – P-glycoprotein [*Schistosoma mansoni*]) docked complex viewed through RasMol 2.1

**Table 9:**
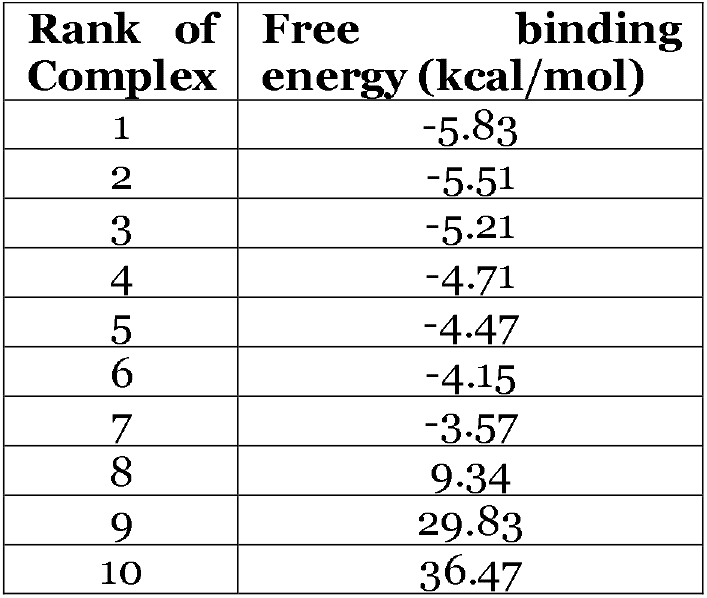
Interaction of the drug Praziquantel with P-glycoprotein [*Schistosoma mansoni*].

**Figure 16:**
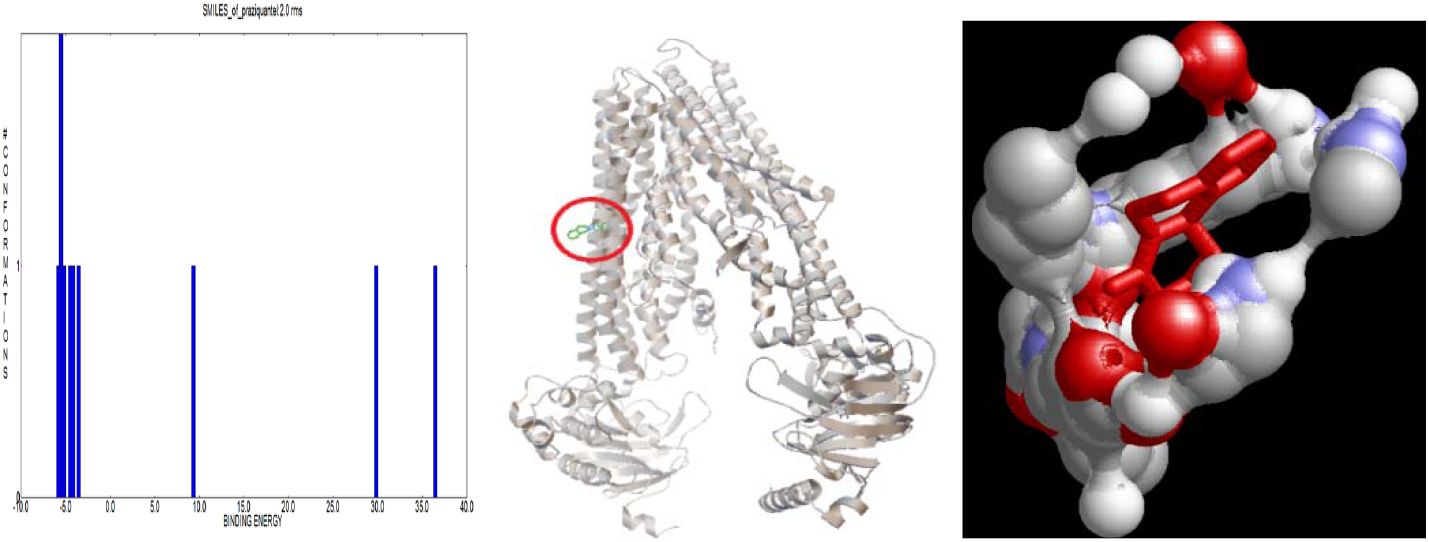
(a) Clustering analysis of the Praziquantel- P-glycoprotein docked complex. (b) Location of the binding site on the receptor (P-glycoprotein [*Schistosoma mansoni*]). (c) The interacting residues in the Praziquantel – P-glycoprotein [*Schistosoma mansoni*]) docked complex is viewed using RasMol 2.1.

#### 3.6.4 Molecular docking results of cladosporin with P-glycoprotein [*Trypanosoma cruzi*]

The best pose has a free binding energy of −6.23 kcal/mol (Table 10). The clustering figure (Figure 17) shows the most number of conformations at −5.0 kcal/mol. Figure 17(b) depicts the binding site on the receptor and figure 17(c) shows the interacting residues in the the cladosporin – P-glycoprotein [*Trypanosoma cruzi*]) docked complex viewed through RasMol 2.1

**Table 10:** Interaction of the drug cladosporin with P-glycoprotein [*Trypanosoma cruzi*]

| Rank of Complex | Free binding energy (kcal/mol) |
| --- | --- |
| 1 | -6.23 |
| 2 | -6.04 |
| 3 | -5.67 |
| 4 | -4.81 |
| 5 | -5.92 |
| 6 | -5.25 |
| 7 | -4.89 |
| 8 | -4.83 |
| 9 | -4.15 |
| 10 | -3.34 |

**Figure 17:**
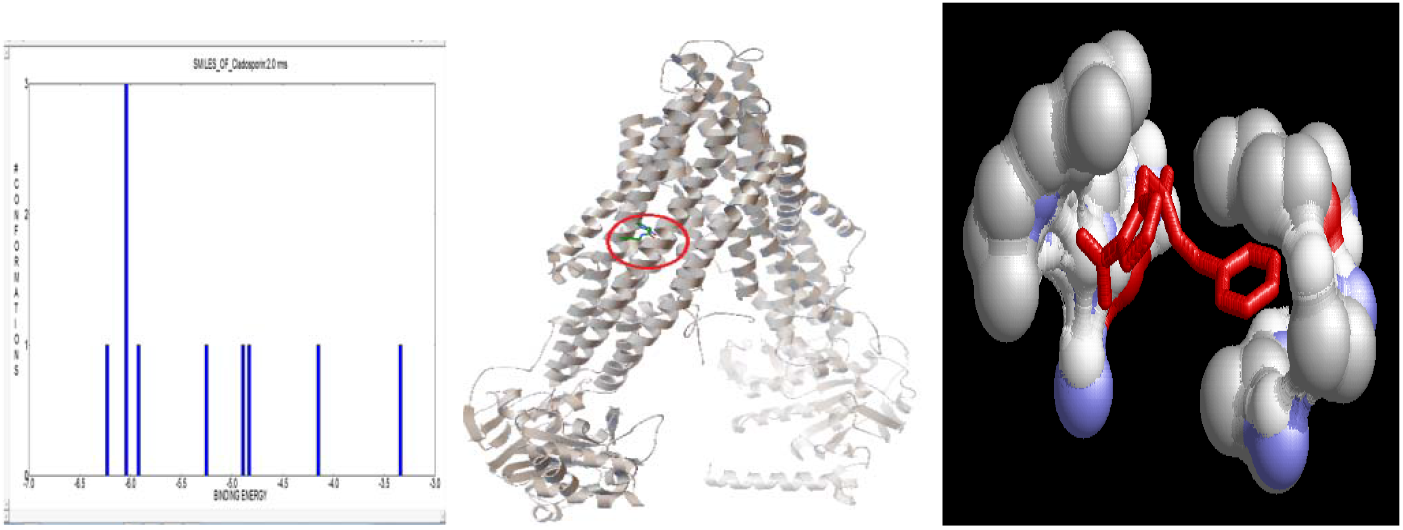
(a) Clustering analysis of the cladosporin-P-glycoprotein docked complex. (b) Location of the binding site on the receptor (P-glycoprotein [*Trypanosoma cruzi*]). (c) cfdaThe interacting residues in the cladosporin – P-glycoprotein [*Trypanosoma cruzi*]) docked complex is viewed using RasMol 2.1.

These steps were carried out for each receptor-ligand complex, and the least free binding energy of each docked complex was determined. These results are summarized in Table 11:

**Table 11:** Free binding energy of all known and investigational drugs, including repurposed antibiotics.

|  | KNOWN DRUGS | FREE BINDING ENERGY | INVESTIGATIONAL DRUGS | FREE BINDING ENERGY |
| --- | --- | --- | --- | --- |
| **L. major** | Amphotericin B | -6.44 | Cladosporin | -6.42 |
|  | Fluconazole | -3.12 | Jaspamide | -5.98 |
|  | Pentamidine | -2.67 | Nifurtimox | -5.66 |
|  | Miltefosine | 1.21 | Praziquantel | -5.59 |
|  |  |  | Dapsone | -5.48 |
|  |  |  | Benznidazole | -5 |
|  |  |  | Tipifarnib | -4.54 |
|  |  |  | Flubendazole | -4.36 |
|  |  |  | Terbinafine | -4.25 |
|  |  |  | Sodium stibogluconate | -3.7 |
|  |  |  | Paromomycin | -3.07 |
|  |  |  | Meglumine antimoniate |  |
| **T. cruzi** | Nifurtimox | -5.22 | Cladosporin | -6.23 |
|  | Benznidazole | -5.02 | Tipifarnib | -5.87 |
|  |  |  | Jaspamide | -5.82 |
|  |  |  | Fexinidazole | -4.62 |
|  |  |  | Suramin | -4.25 |
|  |  |  | Ravuconazole | -3.69 |
|  |  |  | Posaconazole | -2.52 |
|  |  |  | AN2690 |  |
| **O. volulus** | Mebendazole | -5.36 | Praziquantel | -6.16 |
|  | Albendazole | -5.22 | Moxidectin | -5.53 |
|  | Suramin | -4.79 | Niclosamide | -5.29 |
|  | Diethylcarbamazine | -3.76 | Flubendazole | -4.58 |
|  | Ivermectin | -1.35 | Thiabendazole | -4.35 |
|  |  |  | Metrifonate | -2.09 |
|  |  |  | Emodepside | -1.92 |
| **S. mansoni** | Praziquantel | -5.83 | Cladosporin | -6.07 |
|  | Mebendazole | -5.1 | Jaspamide | -6.06 |
|  | Oxamniquine | -4.4 | Niclosamide | -5.72 |
|  | Albendazole | -3.83 | Nifurtimox | -5.62 |
|  |  |  | Artesunate | -5.12 |
|  |  |  | Benznidazole | -4.41 |
|  |  |  | Tipifarnib | -4.38 |
|  |  |  | Imatinib | -4.29 |
|  |  |  | Furozan | -4.27 |
|  |  |  | Suramin | -3.85 |
|  |  |  | Metrifonate | -3.54 |

### 3.7 Calculation of Differential Ligand Binding Affinity

The differential affinity of the potential drug for a given efflux pump protein relative to the known drug is estimated as the difference in the binding energies of the known and potential drugs.

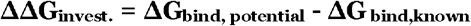

Where ΔΔG_invest_. = Differential ligand affinity, kcal/mol

ΔG_bind_ = Free energy of binding, kcal/mol

For each disease, the differential ligand binding affinity is calculated for every known-potential drug pair. The ΔΔG_investiational_ values are given in Table 12. All values are expressed in kcal/mol. The drugs having ΔΔG_invest_ values greater than the ΔG_invest_. values may have better antihelminthic activity.

**Table 12:**
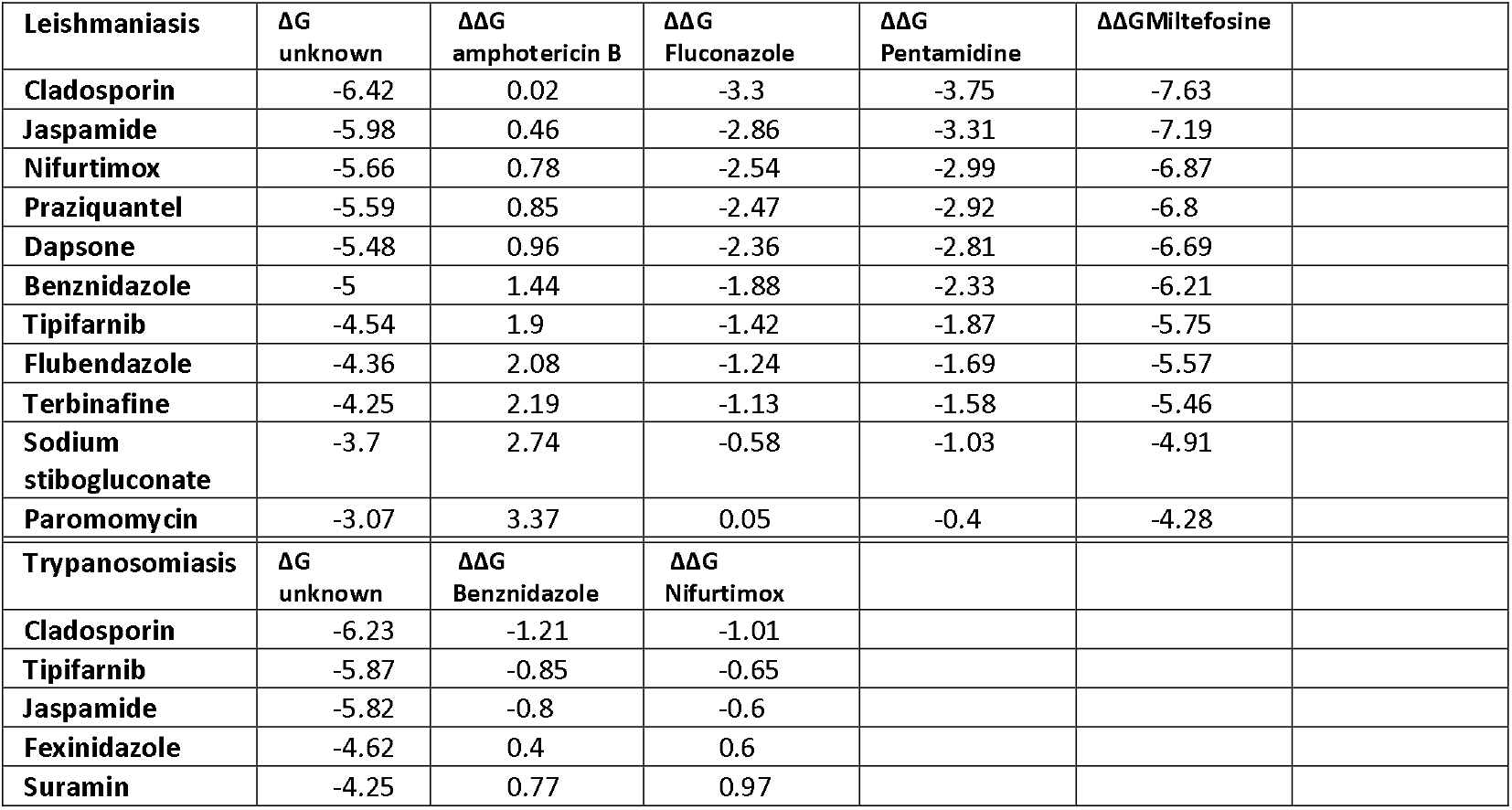
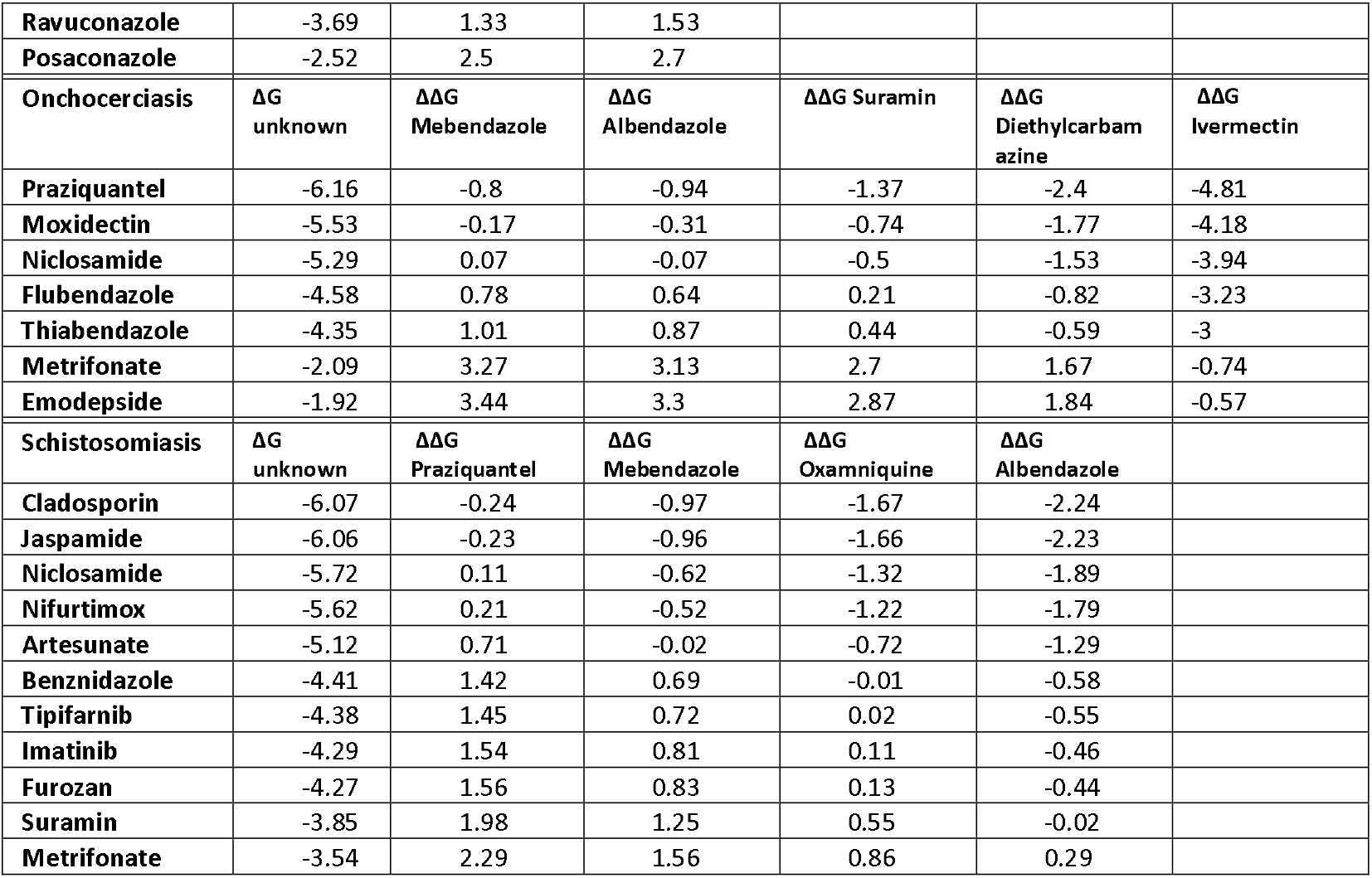
Differential ligand binding affinity for each known-potential drug pair.

All values are expressed in kcal/mol. It can be inferred from these results, that many of the repurposed anti-parasitic drugs show promise for treatment against other helminths. The results shown in table 12 serve as an indicator of which drugs may be promising antihelminthics:

1. Leishmaniasis: Cladosporin (−7.63 kcal/mol), Jaspamide (−7.19 kcal/mol) and Nifurtimox (−6.87 kcal/mol).
2. Trypanosomiasis: Cladosporin (−1.21 kcal/mol) and Tipifarnib (−0.85 kcal/mol)
3. Schistosomiasis: Cladosporin (−2.24 kcal/mol) and Jaspamide (−2.23 kcal/mol)
4. Onchocerciasis: Praziquantel (−4.81 kcal/mol) and Moxidectin (−4.18 kcal/mol)

### 3.8 Analysis of Interacting Residues in each Docked Complex

The best pose of each docked complex was viewed using RasMol 2.1, and all interacting residues within a radius of 4.5 Å of the ligand were restricted and analyzed. The results are summarized in Tables 13-16.

**Table 13:** Interacting residues between the P-glycoprotein of *Leishmania major* and the chosen drugs.

| Receptor | Drug | Interacting residues |
| --- | --- | --- |
| 4M1M | Amphotericin B | Thr172, Asp173, Ser176, Ala683, Asp687, Ser876, Ala879, Leu880, Lys883, Lys884, Glu887, Lys996 |
|  | Fluconazole | Val129, Cys133, Ala136, Asn179, Glu180, Gly181, Gly183, Asp184, Lys185, Met188, Leu875, Asp882, Lys930, Phe934 |
|  | Pentamidine | Glu239, Leu240, Ala242, Tyr243, Ala244, Gly247, Ala248, Glu251, Arg785, Thr811, Ala815, Asn816, Ala819, Gln820 |
|  | Miltefosine | Asp173, Ser176, Lys177, Glu180, Lys185, Leu875, Ala879, Leu880, Lys883 |
|  | Cladosporin | Gln434, Leu437, Leu439, Val468, Ser470, Glu472, Val474, Asn899, Arg901, Thr902, Ser905 |
|  | Jaspamide | Leu254, Ala255, Ala256, Ile257, Arg258, Thr259, Phe800, Asn805, Thr806, Thr807, Gly808, Leu810, Glu1115, Ile1117 |
|  | Nifurtimox | Ala288, Asn292, Gln769, Gly770, Phe773, Gly774, Glu778, Ala819, Gln820, Lys822, Gly823, Ser827, Phe990, Pro992 |
|  | Praziquantel | Leu254, Ala255, Ala256, Ile257, Arg258, Thr259, Phe800, Asn805, Thr806, Thr807, Leu810, Ser1113 |
|  | Dapsone | Val474, Leu475, Phe476, Ala477, Gly521, Glu522, Lys523, Lys891, Thr894, Glu895, Glu898, Asn899, His1003, Arg1006, Ile1007, Lys1010 |
|  | Benznidazole | Asp685, Val688, Pro689, Trp799, Asp802, Lys804, Asn805, Arg813, His1003, Arg1006, Ile1007, Lys1010 |
|  | Tipifarnib | Ala244, Gly247, Ala248, Val249, Glu251, Glu252, Asp1120, Gly1166, Asp1167, Lys1168 |
|  | Flubendazole | Phe159, Asp160, His162, Asp163, Val164, Ser470, Glu472, Val474, Ile897, Gly898, Asn899, Phe900, Arg901, Thr902 |
|  | Terbinafine | Phe159, Val164, Gln434, Gln437, Leu439, Val468, Ser470, Glu472, Val474, Ile897, Glu898, Asn899, Phe900, Arg901, Thr902, Ser905 |
|  | Sodium stibogluconate | Ser470, Glu472, Pro473, Val474, Leu475, Ala477, Glu522, Lys532, Glu895, Glu898, Asn899, Arg901, Thr902 |
|  | Paromomycin | Val164, Glu472, Pro473, Val474, Glu522, Glu898, Asn899 |

**Table 14:** Interacting residues between the P-glycoprotein of *Onchocerca volulus* and the chosen drugs.

| **Receptor** | **Drug** | **Interacting residues** |
| --- | --- | --- |
| 4F4C | Mebendazole | Glu267, Thr268, Tyr271, Ala272, Gly275, Lys276, Lys315, Arg830, Ala860, Thr861, Pro864, Arg867 |
|  | Albendazole | Glu36, Gly37, Asp38, Ile40, Glu267, Thr268, Tyr271, Val305, Ala308, Lys309, Glu823, Thr826, Arg827, Arg830, Ala860, Thr861, Pro864, Arg867 |
|  | Suramin | Lys720, Leu723, Ser724, Lys727, Lys923, Val925, Lys936 |
|  | Diethylcarbamazine | Arg918, Arg919, Phe920, Gly922, Lys923, Asn924, Gln979 |
|  | Ivermectin | Arg172, Thr197, Phe200, Asp201, Glu204, Lys720, Lys923, Asn924, Val925, Ala928, Phe931, Ala932, Gly935, Lys936, Ile939 |
|  | Praziquantel | Leu11, Glu165, Lys207, Asp212, Arg918, Phe920, Lys923, Asn924, Ser927, Phe931, Ala972, Glu975, Gln979 |
|  | Moxidectin | Leu11, Arg12, Asp15, Lys26, Lys30, Glu33, Pro374, Gln913, Tyr914, Arg916, Gly917, Gly1032, Phe1033, Thr1035, Ser1036, Pro1039 |
|  | Niclosamide | Asn4, Gly5, Ser6, Leu7, Ile48, Thr49, Val56, Lys59, Gly380, Thr381, Gln383, Gly384 |
|  | Flubendazole | Glu36, Gly37, Ser42, Thr268, Tyr271, Ala272, Gly275, Arg830, Ala860, Thr861, Pro864, Asn865, Arg867, Lys1043 |
|  | Thiabendazole | Phe504, Asn505, Cys506, Asp933, Lys936, Ile937, Glu940, Phe957, Asn960, Lys964 |
|  | Metrifonate | Gln840, His841, Gly843, Phe844, Ser847, Gln849, Asn850, Lys1057, Ile1058, Lys1060 |
|  | Emodepside | Arg172, Thr197, Phe200, Asp201, Glu204, Asp550, Val925, Ser929, Phe931, Ala932, Gly935, Lys936, Ile939 |

**Table 15:** Interacting residues between the P-glycoprotein of *Schistosoma mansoni* and the chosen drugs.

| **Receptor** | **Drug** | **Interacting residues** |
| --- | --- | --- |
| 4F4C | Nifurtimox | Asn733, Asn734, Gln849, Asn850, Arg1056, Lys1057, Ile1058 |
|  | Benznidazole | Asn4, Gly5, Ser6, Leu7, Thr49, Glu55, Val56, Arg205, Thr381, Gly384, Ala385 |
|  | Cladosporin | Tyr35, Glu36, Ile40, Glu267, Thr268, Tyr271, Ala272, Gly275, Lys315, Arg830, Ala860, Thr861, Arg867 |
|  | Tipifarnib | Gly373, Asp38, Ile40, Asp41, Ser42, |
|  |  | Asn43, Glu267, Thr268, Tyr271, Ala272, Gly275, Arg830, Ser856, Thr857, Ala860, Thr861, Arg867 |
|  | Jaspamide | Gln913, Tyr914, Arg916, Gly917, Arg918, Arg919, Lys923, Gly1032, Phe1033, Thr1035, Ser1036, Phe1038, pro1039 |
|  | Fexinidazole | Gly37, Asp38, Ile40, Asp41, Ser42, Asn43, Thr268, Tyr271, Ala272, Gly275, Arg830, Ser856, Ala860, Thr861, Pro864, Arg867 |
|  | Suramin | ,Lys727, Lys923, Val925 |
|  | Ravuconazole | Ala910, Gln913, Tyr914, Gly917, Arg919, Gly1032, Phe1033, Thr1035, Ser1036, Pro1039 |
|  | Posaconazole | Glu33, Leu161, Gly917, Arg918, Arg919, Phe920, Gly922, Lys923, Asn924, Glu975, Ala976, Gln979, Phe1033, Thr1035, Ser1036, Pro1039 |

**Table 16:** Interacting residues between the P-glycoprotein of *Trypanosoma cruzi* and the chosen drugs.

| Receptor | Drug | Interacting residues |
| --- | --- | --- |
| 4F4C | Praziquantel | Asn505, Arg551, Asp933, Lys936, Ile937, Ile939, Glu940, Glu943, Asn944, Lys964 |
|  | Mebendazole | Arg172, Thr197, Phe200, Asp201, Glu204, Lys207, Asn924, Ala928, Phe931, Ala932, Gly935, Ile939 |
|  | Oxamniquine | Ile40, Asp41, Ser42, Thr268, Tyr271, Ala272, Gly275, Arg830, Ser856, Ala860, Pro864, Arg867 |
|  | Albendazole | Ser42, Glu267, Thr268, Tyr271, Ala272, Gly275, Lys276, Lys315, Arg830, Ala860, Arg867 |
|  | Cladosporin | Tyr35, Glu36, Gly37, Ile40, Phe263, Ala264, Ile265, Glu267, Thr268, Tyr271, Lys315, Arg830, Thr861, Asn865, Arg867, Thr868, Glu1040, Lys1043 |
|  | Jaspamide | Leu161, Lys207, Glu208, Gly211, Asp212, Lys213, Gly917, Arg918, Arg919, Phe920, Gly922, Lys923, Asn924, Gln979 |
|  | Niclosamide | Lys26, Lys30, Ala910, Gln913, Tyr914, Arg916, Gly917, Leu1031, Gly1032, Phe1033, Thr1035 |
|  | Nifurtimox | Tyr35, Glu36, Gly37, Ile40, Phe263, Ala264, Glu267, Thr268, Lys315, Pro864, Asn865, Arg867, Thr868, Glu1040, Lys1043 |
|  | Artesunate | Arg8, Leu11, Arg12, Asp15, Lys26, Lys30, Leu371, Pro374, Arg916, |
|  |  | Phe1033, Thr1035 |
|  | Benznidazole | Asn505, Arg551, Asp933, Lys936, Ile937, Glu940, Lys964 |
|  | Tipifarnib | Leu161, Glu204, Lys207, Glu208, Gly211, Asp212, Lys213, Val378, Asn924 |
|  | Imatinib | Ser42, Asn43, Glu267, Tr268, Tyr271, Ala272, Gly275, Lys276, Arg830, Ala860, Thr861, Arg867, |
|  | Furozan | Ala910, Gln913, Tyr914, Arg916, Gly917, Arg919, Lys923, Gly1032, Thr1035, Ser1036, Phe1038, Pro1039 |
|  | Suramin | Asn4, Arg8, Asp51, Glu55, Thr194, Asp201, Asn202, Arg205, Glu716, Gly719, Lys720, Asp721 |
|  | Metrifonate | Ile40, Phe263, Ala264, Glu267, Thr268, Tyr271, Ala308, Lys315, Arg830, Ala860, Pro864, Arg867 |

The interacting residues are shown and the binding pockets found in each protein sequence with respect to different drugs are highlighted. Analysis of the interacting residues, showed certain binding pockets in each efflux pump protein studied. Certain residues were found to be preferred over others, for drug binding. These preferred binding pockets are:

1. P-glycoprotein [*Leishmania major*]: **(Ser470, Glu472, Val474, Ile897, Glu898, Asn899, Phe900, Arg901, Thr902, Ser905)**
2. P-glycoprotein [*Onchocerca volvulus*]: **(Arg830, Ala860, Thr861, Pro864, Arg867)**
3. P-glycoprotein [*Schistosoma mansoni*]: **(Glu267, Thr268, Tyr271, Ala272, Gly275, Lys276)**
4. P-glycoprotein [*Trypanosoma cruzi*]: **(Arg830, Ala860, Thr861, Arg867)** **(Gly917, Arg918, Arg919, Phe920, Gly922, Lys923)** **(Phe1033, Thr1035, Ser1036, Pro1039)**

## 4. Conclusions

The study of the human P-glycoprotein homologs, namely the P-glycoproteins of *Leishmania major*, *Onchocerca volvulus*, *Schistosoma mansoni* and *Trypanosoma cruzi* has provided an insight into their drug resistance mechanisms. The investigational drugs like Cladosporin, jaspamide, nifurtimox and tipifarnib are strong contenders for novel antihelminthic treatment. Known drugs such as praziquantel and moxidectin have shown great promise for use as treatment against other helminthic diseases

## Notes

https://doi.org/10.6084/m9.figshare.12472091.v1

### Competing Interest Statement

The authors have declared no competing interest.

